# Biotic resistance predictably shifts microbial invasion regimes

**DOI:** 10.1101/2024.10.14.618173

**Authors:** Xiaozhou Ye, Or Shalev, Christoph Ratzke

## Abstract

Invading new territory is a central aspect of the microbial lifestyle, allowing microbes to expand to remote locations and pathogens to spread and infect their hosts. However, invading microbes rarely find novel territories uninhabited. In such a scenario, resident microbes can interact with the newcomers and, in many cases, impede their invasion – an effect known as ‘biotic resistance’. Accordingly, invasions are shaped by the interplay between dispersal and resistance. However, these two factors are difficult to disentangle or manipulate in natural systems, making their interplay difficult to understand. To address this challenge, we tracked microbial invasions in the lab over space and time – first in a model system of two interacting microbes, then in a multi-strain system involving a pathogen invading resident communities. In the presence of biotic resistance, we observed three qualitatively different invasion regimes: ‘consistent’, ‘pulsed’, and ‘pinned’, where, in the third regime, strong biotic resistance stalled the invasion entirely despite ongoing invader dispersal. Surprisingly, these rich invasion dynamics could be qualitatively predicted with a simple, parameter-free framework that ignores individual species interactions, even for rather complex communities. Moreover, we showed that this simple framework could accurately predict simulated invasions from different mechanistic models, indicating its broad applicability. Our work offers a thorough understanding of how biotic resistance impacts invasions and introduces a predictive tool to identify invasion-resistant communities.

## Introduction

Due to their small size, microbes are true masters of dispersal, easily travelling with wind^1,2^, water currents^3^, or other organisms^4^. This dispersal ability allows microbes to reach even the most remote habitats^5^. The dispersing microbes only thrive under the right nutrients^6^, pH^7^, temperature^8^ or salinity conditions^9^. Accordingly, abiotic environmental factors of the novel territory can decide about the establishment of an invading species. However, sequencing data of environmental samples show that microbe species are absent from many habitats that they in principle could live in^10,11^. Are even microbial invasions restricted by limited dispersal, or can also other mechanisms hinder microbial invasion? While dispersal rate (also known as propagule pressure) has been recognized as a key driver for macro-invasions^12–16^, its importance in microbial invasion is inconsistent across studies and therefore remains unclear^17–22^. On the other hand, microbial invasions can also be limited because novel territories are often already occupied by other microbes. These resident microbes can inhibit the establishment of invaders -, an effect known as ‘biotic resistance’ or ‘colonization resistance’^23–26^ - and therefore potentially hinder invasions. The interplay between dispersal and the resident community’s biotic resistance therefore strongly impacts the spatio-temporal dynamics of a microbial invasion (Fig. 1a).

**Figure 1:**
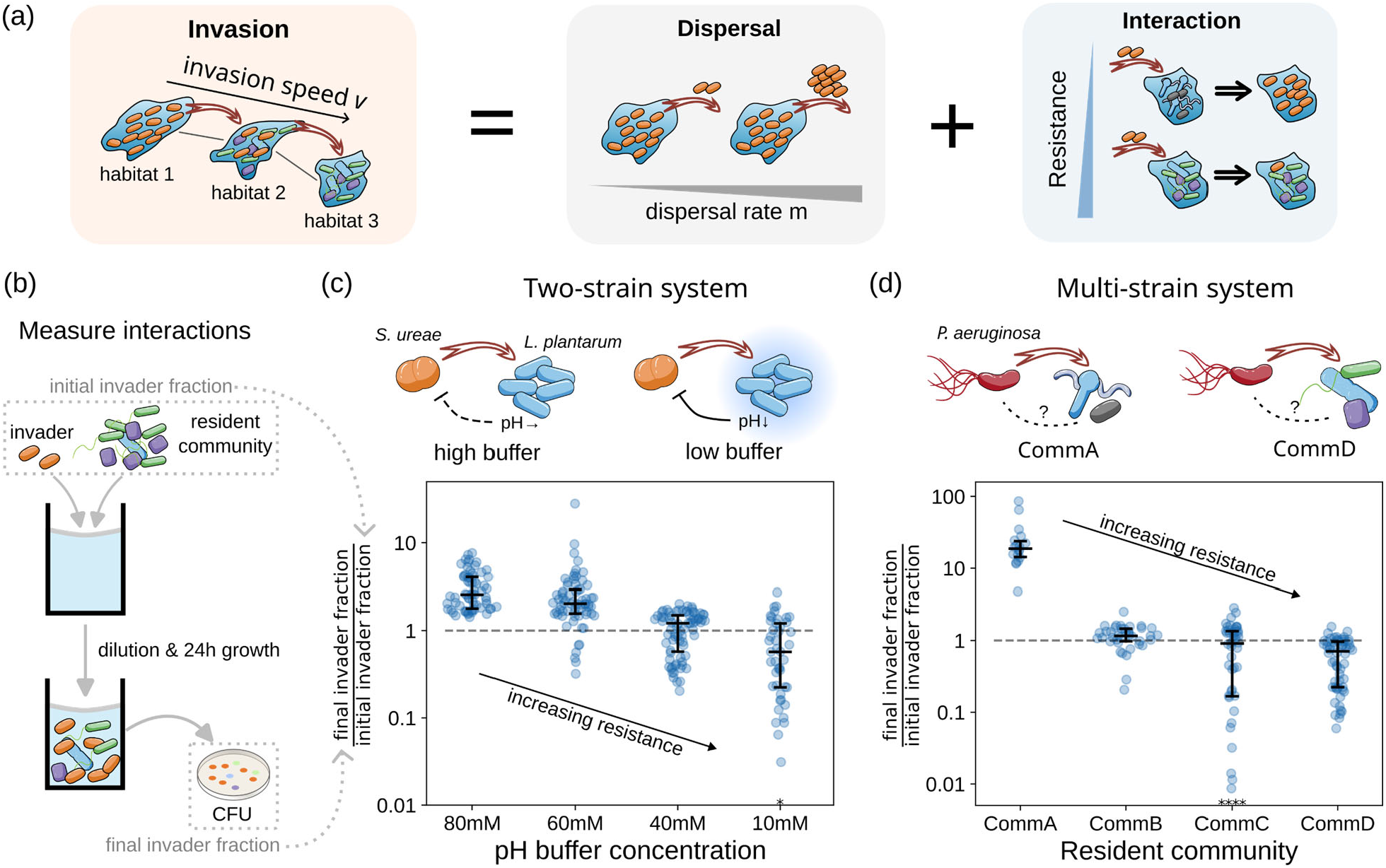
Studying invasions as the combined outcome of dispersal and resistance. **(a)** The dynamics of an invasion (left) depend on the dispersal rate (middle) and the resistance of the resident community (right). **(b)** We measured the interactions between the invader and the resident community to quantify the biotic resistance. We mixed different amounts of an invader to a resident population or community and measured the change of this invader’s fraction after 24h by colony forming units (CFU, see Methods). A lower ratio of final versus initial invader fraction indicates a stronger biotic resistance. **(c)** In a two-strain system with *Sporosarcina ureae* (Su) as invader and *Lactiplantibacillus plantarum* (Lp) as resident species, the biotic resistance of Lp is mainly mediated by pH and can be tuned by changing the media’s buffer concentration. **(d)** In a multi-strain system, four stably coexisting communities of *C. elegans* gut strains show different biotic resistance levels toward the invading pathogen *Pseudomonas aeruginosa* (Pa). Data from different initial invader fractions are plotted if the final invader fraction is < 0.9 and are scattered to reflect probability density distributions. Black horizontal bars denote median and quantiles and black asterisks denote data points that lie outside the plotted areas.

The impact of biotic resistance on invasions can also be relevant for us humans when the invaders are either crop^27,28^ or human^29,30^ pathogens. It is hypothesized that the resident microbiota in our body can inhibit invading pathogens and, in this way, protect us from diseases. In agreement, several studies have shown that beneficial gut microbiota can lower the risk of *C. difficile*^31,32^ and enterobacteria^33–37^ infections. Similarly, in plants, microbial communities of leaves^38–40^ and roots^41–43^ can suppress the growth of pathogens, hence protecting the plant from diseases. Microbial communities that impact diseases within a host may consequently also counteract the spread of infections between hosts. The potential importance of microbial biotic resistance for our personal and global health requires a better and ideally predictive understanding of this phenomenon.

To understand how biotic resistance and dispersal together shape microbial invasions, we must study ecological systems where we can independently manipulate both factors and observe invasions over space and time. In this work, we built microbial systems with different and partially tunable levels of biotic resistance. We used these systems to track invasions in real-time under various dispersal rates and observed that the interplay between dispersal and microbial interactions gives rise to different types of invasion dynamics. The obtained data allowed us to develop a parameter-free framework to predict under which conditions an invasion will be successful. This framework can be used to estimate the invasion resistance of microbial communities, which could, for example, be useful in identifying or even designing pathogen-resistant microbiota.

## Results

### Building microbial systems with varying biotic resistances

To study the impact of biotic resistance on microbial invasions, we first constructed microbial model systems with varying biotic resistances.

As a simple two-species system we chose *Sporosarcina ureae* (Su) as an invader and *Lactiplantibacillus plantarum* (Lp) as resident species. In this system, Lp inhibits the growth of Su by lowering the pH of the media (Supplementary Fig. 1 and previous study^44^), while adding buffer to the media weakens the ability of Lp to change the media’s pH (Supplementary Fig. 1a). Accordingly, the more buffer we add to the media, the less Lp inhibits Su (Fig. 1b,c), weakening the biotic resistance of Lp against Su invasion. In this simple microbial system, we developed the biotic resistance can be thus tuned by the buffer concentration of the media.

To obtain insights into more complex and realistic scenarios of invasion, we also studied the invasion of the pathogen *Pseudomonas aeruginosa* (Pa) into synthetic microbial communities comprising strains obtained from the *Caenorhabditis elegans* gut (the ‘multi-strain system’). For that purpose, we built several resident communities by mixing various sets of strains and letting them settle into their stable states (Supplementary Fig. 2b,c). The obtained communities are composed of different species and differ in their biotic resistances, as can be seen in Fig. 1b, d. These different resistance levels cannot be explained by pH changes alone (Supplementary Fig. 2c). Different from previous studies^45^, the community productivity (OD _600nm_, Supplementary Fig. 2d) also does not determine biotic resistance in our systems. Interestingly, the most resistant community (Comm D) consists of *Pseudomonas* strains (Supplementary Fig. 2b), *i*.*e*. closely related to the invader, suggesting that niche similarity might play a role in determining resistance levels.

The obtained experimental systems offered us a unique opportunity to experimentally investigate the impact of biotic resistance on microbial invasions, as described below.

### Interplay between dispersal and biotic resistance causes three types of invasion dynamics

To observe the impact of biotic resistance on invasion dynamics, we conducted invasion experiments using our two- and multi-strain systems with varying biotic resistances. We performed the experiments along the rows of multiwell plates following a setup used in previous studies^46,47^, starting from 4 wells filled with the invading species, followed by 8 wells with the resident community. Every day we carried out dispersal by transferring a certain fraction (m) from each well to the neighboring wells. Upon dispersal, we also diluted the cultures into a new multiwell plate with fresh media and incubated them for 24h to allow for species interaction (Fig. 2a, Methods). This process was repeated for 10 days. The abundance of the invading species was estimated daily by measuring OD_600_ or bioluminescence (Methods) which correlate with invader fraction for the two- and multi-strain system respectively (Supplementary Fig. 3 and 4). The obtained data allowed us to follow the invasion in space and time and calculate how much distance (wells) the invasion front moved forward both on the n-th day (daily invasion speed v_n_) and on average (mean invasion speed v) (Fig. 2b and Methods). Leveraging this system, we conducted experimental invasion in our simple two-strain system as well as our more complex multi-strain system at varying dispersal rates and biotic resistances. The obtained results are shown in Fig. 2 c-e.

**Figure 2:**
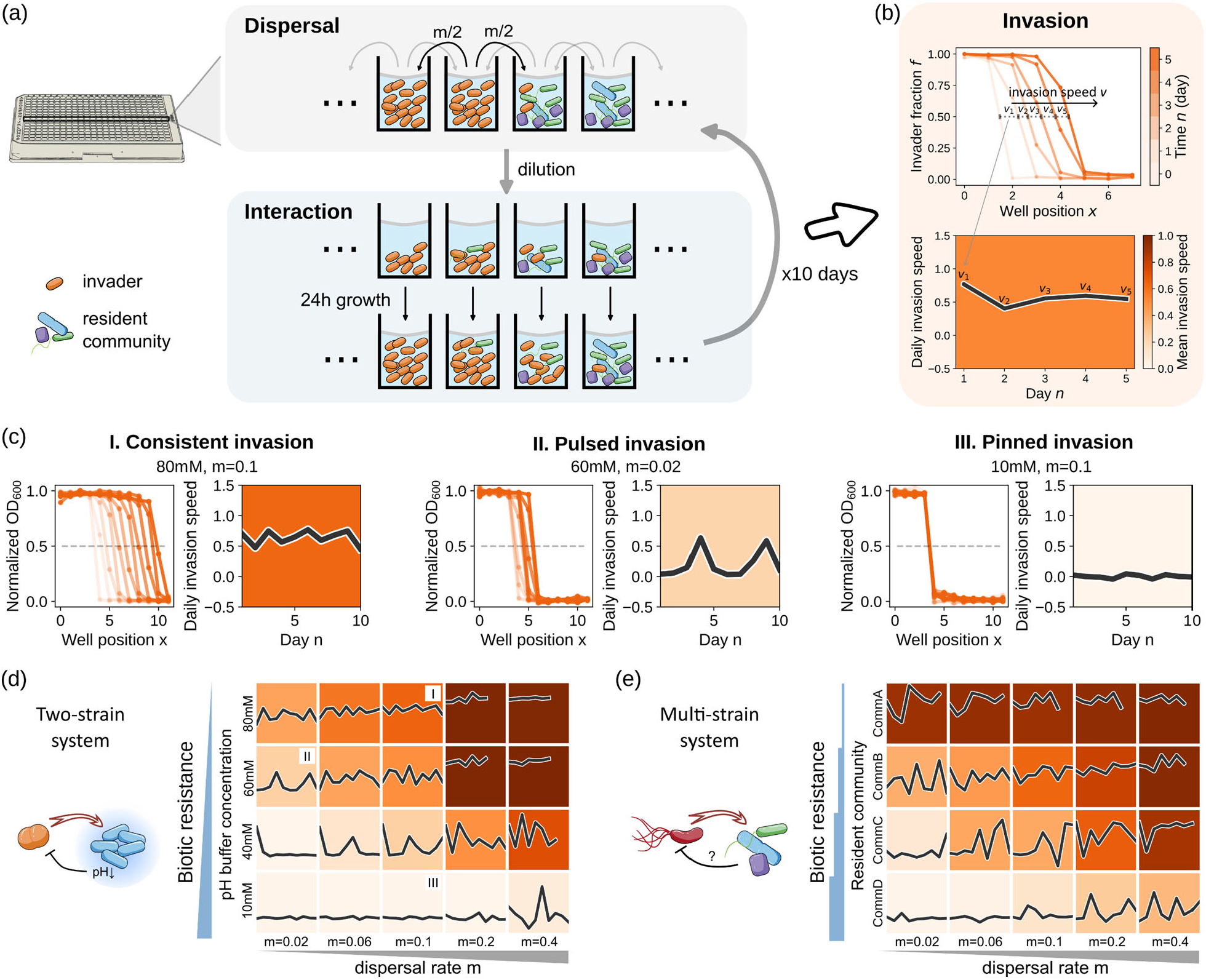
Interplay between dispersal and biotic resistance gives rise to three distinct invasion dynamics. **(a)** We performed invasion experiments in 384-well plates. We carried out dispersal by transferring a fraction m of the bacteria from each well to its neighboring wells. Afterwards, the invader and the resident community interact for 24 hours. We repeated this process for 10 days, as described in more detail in the main text. **(b)** Repeated dispersal and interaction lead to an invasion wave (upper panel), from which the daily invasion speed (black line, lower panel) as well as the mean invasion speed over the duration of the experiment (background color, lower panel) can be obtained. **(c)** Three qualitatively different types of invasions could be observed depending on dispersal rate and biotic resistance: continuous, pulsed and pinned invasions. **(d, e)** Dispersal rate and biotic resistance determine the type of invasion in the two-strain (d) and the multi-strain system (e). The three examples in (c) are labeled by I, II, and III in (d).

**Figure 3:**
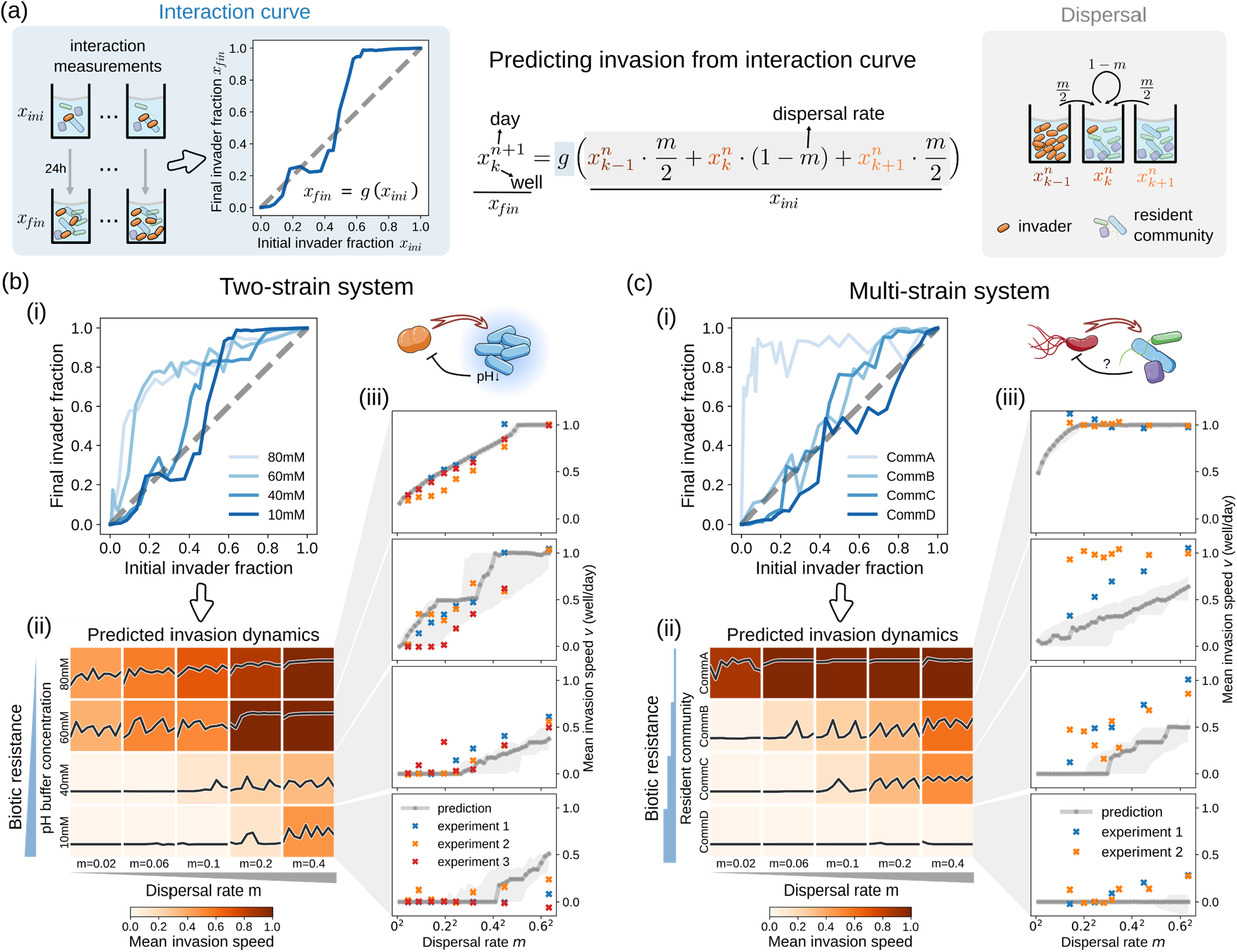
A parameter-free framework predicts invasions in both two and multi-strain systems. **(a)** The parameter-free framework leverages information about biotic resistance (blue background) and dispersal (grey background) to make predictions. The invader fraction in each well on each day is iteratively calculated from the values of the previous day, as outlined by the formula in the middle, which is a combination of Equations 1 and 2 of the main text. We approximate the biotic resistance with an interaction curve g(), which describes the change of the invader fraction after interacting with the resident community. The interaction curve is obtained by mixing the invader and the resident community in different mixing ratios and measuring the invader fraction 24h later (a, left; also shown in Fig. 1b; Methods). Dispersal is represented by a weighted sum of invader fractions in each well and its neighbors (a, right). Since the dispersal rate m is known for each experiment, there are no free parameters left for the model. Details of the model are described in the main text. **(b, c)** Predicted invasion dynamics for two-strain (b) and multi-strain (c) systems. (i): experimentally measured interaction curves for different biotic resistance levels, mean of 3 replicates are shown. (ii): invasion phase diagram predicted from the measured interaction curves using the parameter-free framework depicted in (a); subpanels are as described in Fig. 2, x-axis is day n with range [1, 10], y-axis is daily dispersal rate *v*_*n*_ with range [-0.5, 1.5]. (iii): predicted and measured invasion speeds as dispersal rate changes, for each biotic resistance level. Predictions from the averaged interaction curve (dark grey line) as well as from curves of mean interaction ±1 standard deviation (light grey shadow) are shown. The dispersal rate m is plotted on a squared root scale, which is theoretically predicted to be proportional to invasion speed v in the absence of resistance^48^.

**Figure 4:**
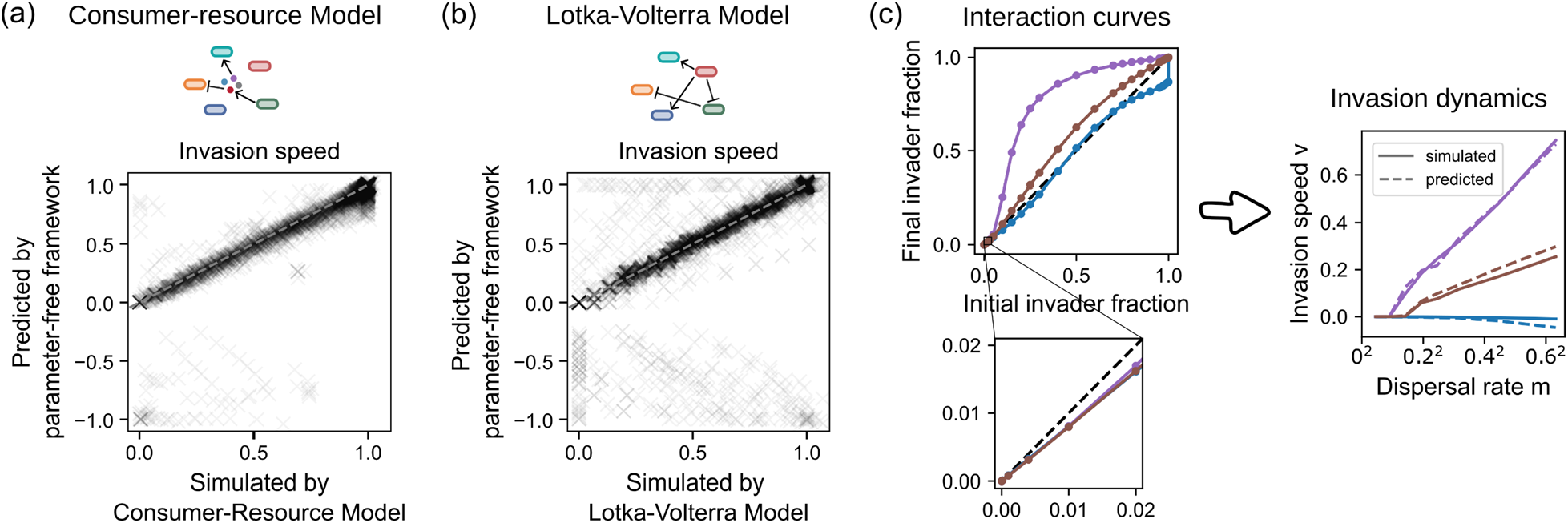
Simulated communities indicate the parameter-free prediction framework is widely applicable. **(a, b)** Predicted vs. simulated invasion speed for invasion simulated by (a) generalized consumer-resource Model where species interactions are mediated by biochemicals (prediction error: median = 0.0071, 90% upper limit = 0.066); and (b) generalized Lotka-Volterra Model where species directly impact the growth of each other (prediction error: median = 0.0059, 90% upper limit = 0.12). For each model, we generated 100 resident communities (each with 4-15 stably coexisting species, Supplementary Fig. 10a, d) and 20 invader species with randomly drawn growth and interaction parameters. For each pair of invader and resident community, interactions curves were obtained from the mechanistic models and inserted into the parameter-free framework depicted in Fig. 3a to make predictions. The invasion simulations were performed equivalent to the experiments shown in Fig. 2a across 8 dispersal rates. Only scenarios with non-negative invasion speed are shown; other scenarios are discussed in Supplementary Fig. 11-14. **(c)** Interaction curves that appear similar at single measurements can be vastly different in overall shape (left), leading to different invasion dynamics (right). Each color corresponds to one invader-resident community pair from the consumer-resource model. The example pairs are chosen such that when initial invader fraction is 0.01 the final invader fraction falls between [0.0079, 0.0081].

We observed three distinct types of invasions: consistent (I), pulsed (II), and pinned (III) (Fig. 2b). In consistent invasion, the invasion front consistently moves forward without a stop (Fig. 2c, I). In pulsed invasion, the invasion front advances in bursts (fast invasion phases) separated by seemingly stationary periods (slow invasion phases) (Fig. 2c, II). During the slow invasion phase, the small invader population faces relatively strong biotic resistance and accumulates slowly at the invasion front (Supplementary Fig. 5e-g); during the fast invasion phase, the invader population growth is large enough to break through the biotic resistance and rapidly take over a new habitat. In pinned invasion, the invasion front is frozen in space and time (Fig. 2c, III). The invader can never overcome the biotic resistance of the resident population, despite ongoing dispersal and consistent arrival of invader cells to the resident community. Accordingly, the invader is permanently present in the invaded habitat patch but can never fully establish growth there even over longer times (Supplementary Fig. 5a-d).

### At strong biotic resistance invaders must overcome a critical dispersal rate to successfully invade

To understand further what determines the type of invasion dynamics, we examined invasions across a wide range of dispersal rates and biotic resistance levels. The resulting phase diagrams are shown for one representative replicate in Fig. 2d-e (for all replicates see Supplementary Fig. 6,7). As can be seen for both the two- and multi-strain systems, consistent invasion can only happen for high dispersal and low biotic resistance. Upon increasing biotic resistance or lowering dispersal, the continuous invasion turns into a pulsed invasion, where the frequency of the pulses gradually decreases and the amplitude increases. Finally, for strong biotic resistance, complete pinning can occur if the dispersal rate falls below a critical value. In other words, resident communities with sufficient biotic resistance can withstand a certain inflow of invaders if it stays below this critical dispersal rate.

Interestingly, this rather simple phase diagram with three distinct regimes of invasion dynamics cannot only be observed for our two-strain system, but also for invasion of *P. aeruginosa* into complex communities, where multiple species interactions determine the outcome. Observing such a simple pattern in such complex systems raises the question if the invasion dynamics could be predicted from simple features of the communities. Such a prediction would allow us to foresee the success of a possible invasion before it happens and identify microbial communities that are particularly resistant towards invasions, which would be of great use in a wide range of fields that deal with microbial invasions.

### Simple interaction measurements predict experimental invasion dynamics

Since an invasion is a combined outcome of dispersal and interaction, understanding both processes well enough should, in principle, allow us to predict the invasion dynamics. However, in practice, we would have to know all interactions within the resident community as well as between the invader and resident species. Such information is, in general, not available. Thus, instead of trying to understand all the mechanistic details of the interactions, we coarse-grained the situation and treated the resident community as “one species” that interacts with the invader. This is a quite strong simplification since most microbial interactions in the system as well as any change in the resident’s community composition during invasion are omitted. The interaction of the invader with the resident community can then be measured by mixing the invader and the resident community in different mixing ratios and measuring the invader fraction 24h later. This shows us the impact of the whole community on the invader, a relationship we termed interaction curve (Fig. 3a, left; Methods). The interaction curve depends on both the interacting species and how these interactions are impacted by the abiotic environment, and thus summarizes both the biotic and abiotic resistance.

Next, we can use the obtained interaction curve to make predictions about invasion dynamics as follows.

Given invader fraction 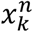 on day *n* and well k *k, k* ∈ 1, 2, … *K*, the updated invader fraction after dispersal (Fig. 3a, right) is given by

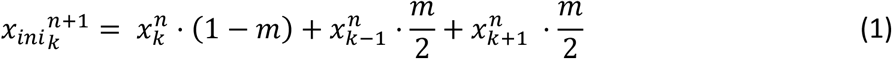

After this simulated dispersal, the change of the invader fraction by species interactions is approximated by the interaction curve as

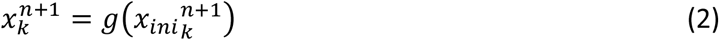

(Fig. 3a, left). Equations 1 and 2 can be combined into a single equation as shown in Fig. 3a, middle. In this way, we obtain a model that does not contain any free parameters and does not explicitly take into account the detailed species interactions in the system. We iterate the above procedure to generate predicted invasion dynamics over days, from dispersal rates and measured interaction curves only.

We next tested whether the proposed prediction framework could predict experimental invasions. To start, we measured the interaction curves as described in Fig. 3a, for our two-strain (Fig. 3b(i), Supplementary Fig. 8) and multi-strain systems (Fig. 3c(i), Supplementary Fig. 9). Then, we inserted the interaction curves into the prediction framework in Fig. 3a to predict invasion dynamics. It is important to note that we measured the interaction curves in a separate experiment and did not obtain them from the invasion data. Furthermore, the points on the interaction curve are not correlated with each other as we measured them from independent measurements, i.e. different bacterial cultures. For the two-strain system, the predictions not only capture the qualitative shift of invasion dynamics as resistance and dispersal rate change but are even quantitatively accurate (Fig. 3b). For the multi-strain system, the predictions still qualitatively capture the invasion dynamics and its transition from pinned to pulsed to consistent as dispersal rate increases and biotic resistance decreases (Fig. 3c). Our prediction however seems to underestimate invasion speed, possibly due to noise in experiments which has been theoretically shown to mostly accelerate invasion^49^.

Interestingly, the observed interaction curves have rather complex shapes, showing that the residents’ ability to resist the invader frequently varies depending on the relative abundance of the invader. Indeed, the overall shape of these interaction curves determines the invasion outcome more than the interaction at a fixed invader density, as discussed in more detail in the next section. The shape is especially important when the interaction curve crosses the diagonal. If the resident community can reduce invader fraction at low initial invader fractions but not at high initial invader fractions (e.g. 40mM and 10mM buffer in Fig. 3b; communities C and D in Fig. 3c) the dispersal rate becomes important for determining the invasion outcome, causing the emergence of a critical dispersal rate (Fig. 3b(iii), c(iii)).

The success of our simple prediction framework means that for a given invasion scenario, we could simply measure the interaction curve to get rather good answers to whether and how invasions would proceed under different dispersal scenarios, without knowing the mechanistic details of the interactions. Measuring an interaction curve is much faster and easier than tracking an invasion over time, allowing us to predict invasions before they even occur. Moreover, since the interaction curve isolates the interaction component from the invasion process, it can be applied to different dispersal rates and invasion scenarios.

### Simulated invasions indicate the prediction framework is widely applicable

To understand how generally the parameter-free framework of Fig. 3a can predict invasion dynamics, we used it to predict invasions simulated from two mechanistic models: the generalized consumer-resource model and the generalized Lotka-Volterra model. We first generated “ground truth” data with the mechanistic models as described in the Methods and Fig.4 captions and then investigated how far the parameter-free framework can recapitulate these data (Fig. 4 a,b). The invasion speeds predicted by the parameter-free framework correspond surprisingly well with the results of both mechanistic models, even for rather sbiodiverse resident communities (Supplementary Fig. 10 b, e). In most cases, the invasion speed fluctuations are also predicted correctly (Supplementary Fig. 10c, f). Prediction failure happens only rarely and often for more complex invasion dynamics like backwards invasion and bidirectional invasion (Supplementary Fig. 12). The framework can also predict invasion in other dispersal scenarios, for instance when dispersal is unidirectional instead of bidirectional (Supplementary Fig. 13).

Such high prediction accuracy is surprising, because contrary to the mechanistic models our parameter-free framework neglects most microbial interactions as discussed above. Moreover, the interaction curve that describes the impact of the resident community on the invader is assumed constant and thus unaffected by the invader (Fig. 3). But in reality, we expect that the invader usually alters the composition of the resident community and thus in return how the community impacts the invader. We suspect that the parameter-free framework is robust against this strong assumption because the success of an invasion is mostly decided in its early phases where the density of the invader is low and its impact on the resident community only minor. This statement is underlined by a closeup comparison of the predicted and simulated invasion dynamics that are discussed in more detail in Supplementary Fig. 11.

We further found that the predictions of our parameter-free model depend mostly on the shape of the interaction curve and less on the interaction outcome for a specific initial invader fraction. As a demonstration, Fig. 4c shows three interaction curves taken from the consumer-resource model, each reaching a similar final invader fraction upon starting with 1% invader (Fig. 4c, left, insert), whereas the overall shapes of the curves are different (Fig. 4c, left). Despite their similarity at small initial invader fractions, the different curve-shapes result in different invasion dynamics (Fig. 4c, right). Therefore, it may not be sufficient to study invasions at a fixed invader fraction. We provide a more intuitive understanding of the connection between shape of the interaction curve and resulting invasion process in the supplement by representative examples (Supplementary Fig. 14) and by a graphical approach called cobwebbing (Supplementary Fig. 15). Importantly, approximating the interaction curve with 3-5 datapoints is normally sufficient for accurate predictions (Supplementary Fig.16). This high data-efficiency makes our parameter-free framework applicable even for invasions with limited data available. Overall, our results suggest that interaction curves summarize the key aspects of biotic resistance and can be used to predict invasion in a wide range of practical contexts.

## Discussion

Microbes usually intrude into space that is already occupied by resident organisms. Consequently, the invasion outcome not only depends on how fast an invader can spread into a novel territory and the abiotic conditions of the new habitat but also on how it interacts with the resident organisms at the new location. Many current studies focus on one of the two aspects: either the spatial expansion of microbes into empty space^47,50,51^ or the interactions between microbial invaders and residents without spatial context^22,52–56^. Our study extends these works by showing that complex yet predictable dynamics of invasions can emerge from the interplay between both dispersal and biotic resistance as caused by species interactions.

Upon changing dispersal and biotic resistance, we observed three types of invasion dynamics in our experiments: consistent, pulsed, or pinned invasion waves. At weak biotic resistance, consistent invasion occurred, just like a classical invasion into empty space^48^. As biotic resistance increased, and the dispersal rate decreased, pulsed and pinned invasion waves emerged. Theoretical studies modeled invasion as a single species reaction-diffusion process and identified two conditions for pulsed and pinned waves. The first condition is patchy instead of continuous habitats^57–59^, as is the case for our experiment and many natural scenarios^60^. The second condition is Allee effect^57,61–63^, which means the growth rate of a species reduces when the population size drops, often due to the need for cooperative growth^64^. In our study, biotic resistance of resident communities makes it harder for the invader to grow at low population density as compared to high density, leading to similar dynamics as caused by an Allee effect. Although we performed dispersal as a discrete daily event, pinned and pulsed invasion can also happen under continuous dispersal^57,58^. Our results indicate that simplified theories about the spread of a single species can be applied to more complex scenarios where resident communities are involved.

The occurrence of pulsed and pinned invasion can have important ecological consequences. During a pulsed invasion, the invasion seemingly comes to a stop at times just to accelerate later again. Such dynamics could be misinterpreted as a halt of invasion when the observation period is too short. Pulsed invasion dynamics have been observed in macro-ecosystems^65^ and were explained by stratified dispersal and Allee effect of the invader. Our findings indicate that such dynamics can also be related to biotic resistance. Since natural habitats are usually already inhabited, we expect that biotic resistance to be a very common effect in nature. Pulsed waves are only mildly affected by spatial heterogeneity in dispersal rate (Supplementary Fig. 17) and carrying capacity (Supplementary Fig. 18); heterogeneity in growth rates has a bigger impact and can frequently lead to irregular pulses (Supplementary Fig. 19). At strong biotic resistance and low dispersal, pulsed invasion turns into pinned invasion, in which the invasion front is completely frozen in space and time. Despite ongoing dispersal, the pinned wave forms a stable boundary between the invader and the residents due to microbial interactions. Such boundary formation may explain some macroscopic patterns, such as the alternating ^66^.

Biological invasions can cause major damage to ecosystems and human society^29,30,67–70^. Accordingly, strong efforts are undertaken to stop invading species around the world. We show here that biotic resistance is crucial to support these efforts. Strong biotic resistance makes pinned and, therefore, a complete stop of an invasion possible, if the dispersal rate is below a critical value (i.e., critical dispersal rate). Without a critical dispersal rate, the invasion will progress even with a very low dispersal rate. Consequently, the invader must be completely obliterated to stop an invasion which might be very challenging in practice, making biotic resistance almost a precondition for stopping invasions.

Identifying potentially invasive species and fragile resident communities before invasion happens is a big challenge in ecology. Several approaches have been proposed to spot invasion risk based on traits of potential invaders^71–75^ or properties of resident communities such as biodiversity^22,56,76–81^ but with mixed success. Our parameter-free framework takes an alternative approach; instead of focusing on specific properties of potential invaders and resident communities, it bases predictions on coarse-grained measurements of the interaction between the two (i.e., interaction curve). For a given invader and a corresponding resident community, the framework predicts how fast invasion would proceed under different dispersal rates, particularly if a critical dispersal rate exists. This framework allows us to identify invaders that would easily invade a given ecosystem and to estimate how much a given community can resist an invasion. Our approach is, in principle, applicable across various ecosystems. It surely comes with the limitation that measuring the biotic resistance and dispersal rate can be difficult for many macroscopic ecosystems. Nevertheless, such measurements are generally feasible in microbial ecosystems, and hence our approach may be applied to estimate the risk of disease spreading in the human gut or skin for a given microbiome profile.

Having a reliable indicator for biotic resistance can also be useful for manipulating, or even rationally designing microbial communities towards higher resistance. Many studies aim to understand how environmental factors impact the resistance of microbiota, such as drugs^82,83^ and diet^84,85^ in the human gut or light conditions in plants^86^. Moreover, teams from both academia^87,88^ and industry^89–91^ are trying to build resistant microbial communities from scratch. For both applications, it is essential to estimate the pathogen resistance of microbial communities. This is normally done by introducing a certain amount of pathogen into the microbiota and observing its growth, which may not capture all features of biotic resistance as we discussed in Fig. 4c. We extend this approach by using slightly more measurements to obtain the interaction curve and, through the prediction framework, linking it directly to predicted invasion outcomes. Importantly, the framework predicts invasion for any given dispersal rate, thus providing a more comprehensive definition of biotic resistance.

The other way around, microbiota engineering can also require a successful invasion. For instance, gut microbiota are transferred from healthy to sick people (i.e., fecal transplantation)^32,92,93^ to treat gut-related diseases, or specific bacteria strains are consumed (i.e. probiotics) to restore healthy gut microbiota^94–98^. Similarly, there are attempts to fight plant pathogens by introducing protective bacteria^99–101^, or to improve productivity by adding fertilizing microbes^102,103^. In all cases, the introduced microbes have to invade the resident microbiota to fulfill their task, which can often be challenging^104,105^. Our framework could predict invasion outcomes for different doses and frequencies of microbe treatment (i.e., different dispersal) and help optimize therapies.

Specifically, if a critical dispersal rate is predicted, a sufficient number of microbes must be added to overcome it.

Taken together, we have shown that biological invasion into occupied territory results in a rather simple set of invasion dynamics, despite complex species interactions and spatial processes. These dynamics can moreover be predicted by a straightforward, parameter-free framework, suggesting that it is feasible to understand and manipulate invasion for our benefit.

## Methods

### Preparation of media and agar plates

Bacteria were precultured in nutrient media (NM), consisting of 1% yeast extract (Sigma-Aldrich #70161) and 1% soytone (Peptone from soybean, Sigma-Aldrich #87972). Both interaction and invasion experiments were performed in invasion media (IM), which resembles diluted De Man, Rogosa, and Sharpe Media (MRS) with some compounds substituted by more commonly used chemicals in our lab. IM consists of 0.1% soytone, 0.1% tryptone (Sigma-Aldrich #95039), 0.04% yeast extract, 7.34mM ammonia acetate (Merck #1.01115.1000), 1.64mM tri-sodium citrate (VWR #27833.237), 0.8mM MgSO_4_, 0.2mM MnSO_4_, 0.4% glucose, 12mM (NH_4_)Cl, 12mM NaCl. Different concentrations of phosphate buffer (pH=6.8, made from 0.58M K_2_HPO_4_ and 0.42M KH_2_PO_4_ followed by pH adjustment) were added to IM, as specified for each condition in the two-strain system and fixed to 10mM for the multi-strain system. Before each experiment, IM was freshly assembled from 6 stocks with partial components (Supplementary Table 1), sterilized by 0.2um filter (Filtropur S, Sarstedt #83.1826.001), and stored in 4C for up to 2 weeks. Such assembly from partial stocks reduced interactions between components during storage and enhanced reproducibility.

Colony forming units (CFU) of bacteria cultures were quantified on different varieties of NM agar plates depending on the experiment. NM agar plates of pH 5 and pH 9 were prepared from 1% yeast extract, 1% soytone, 10mM NaH_2_PO_4_, and 2.5% agar (Agar-Agar, Carl Roth #5210.2), pH was adjusted before autoclave sterilization. pH7 NM agar plates with/without gentamicin were prepared from 1% yeast extract, 1% soytone, and 2.5% agar, for plates with antibiotics, 5µg/mL gentamicin (Gentamicin sulfate, Acros Organics #455310050) was added after the agar partially cooled down following autoclave sterilization. 45mL autoclaved agar mixture was poured into a 150mm petri dish for each agar plate.

### Luciferase tagging of the invader in the multi-strain system

*Pseudomonas aeruginosa PA01* (‘PA01’) was genomically transformed with the luxCDABE operon using the mini-Tn7 insertion system as previously described^106^. Briefly, the plasmids pUC18T-mini-Tn7T-Gm-lux (Addgene #64953) and pTNS2 (Addgene #64968) were curated from E.coli DH5a using a plasmid extraction kit (peqGOLD Plasmid MiniPrep Kit II; VWR, Leuven, Belgium). PA01 cells were made competent for transformation by treatment with a sucrose 300 mM solution PA01, and were then electroporated with 50 ng of the curated pUC18T-mini-Tn7T-Gm-lux and pTNS2 plasmids. After recovery in 1 ml of LB at 30 °C, the electroporated cells were plated on an LB-agar+Gm30 plate. Plates were incubated for 24 hours at 30 °C, and colonies were tested for their bioluminescence activity using a plate reader. Positive colonies were then further grown in a selective medium (LB+Gm30), and were stocked in 25% glycerol at -80 °C.

### OD_600_ and bioluminescence measurement

#### OD_600_ (two-strain system)

A 384-well plate with bacteria cultures was mixed for 15 seconds at 2000rpm on a Mixmate Plate Shaker (‘Mixmate’, Eppendorf, Hamburg, hereon shortened as Mixmate), then put into FLUOstar Omega Plate Reader (BMG Labtech, hereon shortened as plate reader) to measure absorbance at 600nm.

#### Bioluminescence (multi-strain system)

The bacteria cultures were transferred to a Lumox 384-well plate (Sarstedt 94.6130.384) for bioluminescence measurement. First, 2.5µL NM was dispensed into the Lumox plate, then 20µL bacteria culture was reverse dispensed into the Lumox plate using Viaflo384 (Integra) and mixed with the NM for 30s at 1500rpm in Mixmate. The Lumox plate was incubated at 30C in a plate reader for 2min, allowing the added NM to reactivate potentially stagnated bacteria (Pa) metabolism, which gave rise to a more reliable bioluminescence signal. Finally, bioluminescence was measured with a luminescence emission filter, gain 3600, and exposure time 1s.

The raw bioluminescence measurement was corrected for signal crosstalk and background using an empirically fitted function. For measured bioluminescence value *x*_*j*_ of row i and column j, the corrected value 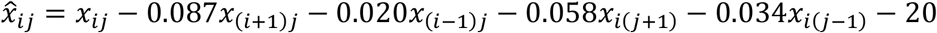. Before correction, signal crosstalk can lead to up to +1800 (a.u.) measurement error, which was reduced to within ±200 (a.u.) after correction.

### Community composition based on colony forming unit (CFU)

In interaction experiments as well as representative invasion experiments, we obtained species composition of bacteria cultures by CFU. First, we serially diluted cultures 1/10 in PBS until reaching a dilution factor of 1/10^6^, the dilutions were carried out in a plate-to-plate manner. Then, for each 96-well intermittent pattern in the 384-well plate, we used Viaflo384 to reverse pipette 5µL from each dilution onto different varieties of NM agar plates, getting 96 separate droplets on the agar surface. For the two-strain system, we used pH 5 agar plates for selective plating of Lp and pH 9 agar plates for Su. For the multi-strain system, we used pH 7 agar plates with gentamicin for selective plating of Pa and pH 7 agar plates without gentamicin for both the resident community and Pa, on which colonies of Pa could be morphologically identified under stereoscope. After the droplets dried out, we incubated the agar plates at 30C for 24 hours until colonies grew to a distinguishable size. We then either took photos of the plates under ring light illumination and counted CFU afterward (for the two-strain system) or directly counted CFU under the stereoscope (for the multi-strain system).

### Community composition based on 16S sequencing

For the multi-strain system, the resident communities’ compositions were confirmed by 16S sequencing (Supplementary Fig. 2 b, c). We extracted DNA from stable cultures of resident communities using peqGOLD Bacterial DNA Kit (VWR #13-3450-02), then sent the obtained DNA to Eurofins for library preparation and sequencing of 16S rDNA V3-V4 region. To determine species composition, we aligned the obtained reads against a custom library comprising the V3-V4 region of the 16 species that we used, which were previously obtained by Sanger sequencing^107^. We carried out alignment using Bowtie2 in Python under ‘--very-sensitive-local’ setting.

### Assembly of stable resident communities in the multi-strain system

#### Summary

As shown in Supplementary Fig.2, we randomly assembled 384 communities each from a subset of a pool of 16 bacteria strains. We cultured the resulting community in 50µL IM (10mM buffer) on a 384-well Labcyte plate (Beckman #001-1455) for 8 days, with 1/200X daily dilution. The 384 initial communities collapsed into ∼5 different representative compositions as indicated by CFU morphologies. We picked 4 wells with distinct compositions and stocked in 30% glycerol at -80°C. The frozen stocks were revived and cultured for 4 additional days with 1/200X daily dilution before being used for experiments. More details are as follows.

#### Preculture and handling of single strains

We assembled the resident communities used in the multi-strain system from 16 strains that were previously isolated from *C. elegans* gut^108^. The strains cover diverse phylogeny (Supplementary Fig. 2b) and have distinct colony morphologies on NM agar plates. We streaked out the 16 strains on pH 7 NM agar plates, then took a single colony from each strain and precultured in 200µL TSB (Tryptic soy broth non-animal origin irradiated, VWR #84674.0500) in a 96-well 500µL deep plate (Masterblock 96well, Greiner #786201). We sealed the plate with 2 layers of Breathable Rayon Films (VWR #391-1261, hereon shortened as breathable films), then incubated on Titramax 1000 shaker (Heidolph) at 1350rpm and 30°C overnight (16h). We then washed precultures twice and resuspended each in 200µL IM (10mM buffer).

#### Assembly and daily dilution

We transferred 50µL of washed bacteria preculture to a 384-well Labcyte plate, as the source plate for community assembly. We added 50uL/well IM (10mM buffer) to another 384-well Labcyte plate as the destination plate. For each well in the destination plate, we randomly assigned the presence (p=0.75) or absence (p=0.25) for each species, then transferred from source plate 25nL of each species that was assigned presence for the current well. This resulted in 384 different initial communities consisting of 0-16 species (mean = 12), and 2 technical replicates with identical community layouts. We sealed the destination plate with 2 layers of breathable films, incubated on Titramax 1000 shaker at 1350rpm and 30°C for 24h. We continued culturing the communities for 7 consecutive days with 1/200X daily dilution, by transferring 250nL into a new 384-well Labcyte plate with 50uL/well fresh IM (10mM buffer).

#### Freezing, reviving, and preparing glycerol stocks of the communities

On day 8, we mixed 25µL of each resulting community with 25µL 60% glycerol and froze the plate at -80°C. To observe community compositions, we diluted the communities with 1/10x dilution series and droplet plated 5µL of the 1e-7, 1e-6, and 1e-5 dilutions using Viaflo384 onto pH7 NM agar plate. The 384 communities collapsed into ∼5 distinct compositions, as indicated by plated CFU morphologies (Supplementary Fig. 2b). We selected 4 communities with distinct compositions, thawed the frozen cultures, and cultivated them in 384-well Labcyte plate and 50µL IM (10mM buffer), with 1/200X daily dilutions for 8 more days to stabilize the communities. We froze the resulting cultures in 30% glycerol at -80°C. We revived resident communities from these stocks by 1/200X daily dilution into 384-well Labcyte plate with 50µL IM (10mM buffer) and culturing on Titramax 1000 shaker at 1350rpm and 30°C in between. We confirmed by CFU and 16S sequencing that community compositions reached equilibrium within 3 days (Supplementary Fig. 2b). We performed all multi-strain system experiments from these stocks.

### Invasion experiments

#### Overview

We carried out invasion for each given resistance level and dispersal rate m in 12 wells in a 384-well plate, mimicking 12 habitat patches. We cultured the invader and the resident community separately for 3 days to stabilize the potential initial dynamics, then initialized the invasion experiment with 4 wells of invader followed by 8 wells of resident community. Every day during the invasion experiment, we performed dilution and dispersal from the current plate to a new plate with fresh media. Specifically, given the dilution rate δ (1/200X unless otherwise specified) and dispersal rate m (8 different values, as specified below), we diluted δ· *m* of the total volume from each well to the same well in the new plate, and 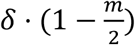 of the total volume to its left and right neighboring wells respectively (or the same well if no left/right neighbor) in the new plate. Essentially, we performed a bidirectional dispersal between adjacent wells. After dispersal and dilution into fresh media, we cultured the new plate and let the species interact for 24 hours (see below for study system-specific culture conditions). We recorded the spatial distribution of the invader daily by measuring OD600 for the two-strain system/bioluminescence for the multi-strain system, and for one experiment (Supplementary Fig. 5) also community composition by counting distinct colony forming units (CFU). We repeated this dispersal-interaction cycle for 8-10 days. We performed 3 biological replicates for the two-strain system and 2 biological replicates for the multi-strain system, the replicates were run independently and in different months.

#### Two-strain system

We streaked out Su and Lp from glycerol stocks to pH 7 NM agar plates and grew them at 30°C for a day. We picked a single colony from each species with an incubation loop and cultured it in 5mL NM in a falcon tube with the lid fully closed to avoid cross-contamination (the species still grew), on Innova 2000 shaker (New Brunswick, Eppendorf AG) at 225rpm and 30°C overnight (∼16h). We took 400µL from the preculture for each species, washed the sample twice, and resuspended the cells in 400µL IM (without buffer). We transferred 250nL of the resulting cell suspension into 384-well Labcyte plates that contained 50µL IM with corresponding concentrations of phosphate buffer. We sealed the plates with two layers of breathable films and incubated them on Mixmate at 1500rpm and 30°C for 24h. We cultivated the bacteria for 3 more days with 1/200X daily dilution to eliminate memory from the preculture and to stabilize the potential population dynamics before starting the invasion experiment. The daily dilution was performed by transferring 250nL from previous plate to a new plate with 50µL IM with corresponding buffer concentrations.

We performed invasion experiments in Labcyte 384-well plates with 50µL IM with corresponding buffer concentrations. We used each one of the four intermittent 96-well patterns in the 384-well plate to perform invasion experiments for one buffer concentration. The invasion was carried along each row (12 wells) of the 96-well pattern, every row corresponded to a different dispersal rate m (m= [0.002, 0.008, 0.02, 0.038, 0.06, 0.1, 0.2, 0.4]). The experiments consisted of dispersal and 1/200X daily dilution, followed by 24h growth on Mixmate at 1500rpm and 30°C. Every 24h, we started the transfer by mixing the cultured plate for 15s at 2000rpm with the Mixmate, then diluted 250nL into another 384-well Labcyte plate with 25µL fresh IM. From the original plate (1X) and the diluted plate (1/100X), we performed dispersal and dilution into a new 384-well Labcyte plate as specified in Supplementary Table 2. After the transfers, we sealed the plates with two layers of breathable films and incubated them in the Mixmate at 1500rpm and 30°C for 1 day (slightly less than 24h, due to the time needed for performing dilution and dispersal). The original plate (1X) was used to measure OD_600_, and in cases where community composition was needed, the diluted plate (1/100X) was serially diluted and plated for CFU analysis (Methods).

#### Multi-strain system

The experiment followed the same principle as the Su invading into Lp experiment, with a few modifications due to the technical difficulties regarding reliably transferring Pa and the communities, as specified below.

Modified culturing media. Instead of using IM with different phosphate buffer concentrations, the phosphate concentration was fixed to 10mM.

Modified preculture procedure. We did the first preculture of Pa in 5mL NM similarly to Su/Lp. We did the first preculture of the community by 1/40 dilution from glycerol stock to 50µL IM (10mM buffer) in 384-well Labcyte plate. We cultured the plate on a Heidolph Titramax 1000 shaker at 1350rpm and 30°C for 24h. After the first preculture, we cultured both Pa and the communities in a 384-well Labcyte plate for 3 more days with 1/200X daily dilution to stabilize possible initial community dynamics. We carried out the dilutions with the tip-based Viaflo384 pipetting system.

Modified liquid transfer during the invasion. Compared to the Su/Lp invasion experiments described above, we slightly modified dispersal rates (m= [0.02, 0.04, 0.06, 0.08, 0.1, 0.12, 0.22, 0.4]) to use only 1/10X diluted bacterial culture in source plates. This modification was made because 1/100X could not be reliably transferred because Pa tended to gather on the surface. Every day, we used Viaflo384 to dilute 5µL bacteria culture into a 384-well Labcyte plate with 45µL IM (10mM buffer). We mixed the 1/10X diluted plate 15s at 2000rpm with Mixmate, let rest for 2min, then performed dispersal and dilution as specified in Supplementary Table 3. We used each one of the four intermittent 96-well patterns in 384-well plate to perform invasion experiments with one resident community. After liquid transfer for dispersal and dilution, we cultured the plate on Titramax 1000 shaker at 1350rpm and 30°C for slightly less than 24h before performing the next dispersal.

#### Invasion speed

We defined the invasion front as the furthest well where the invader population crossed the 50% threshold, i.e. the midpoint of the invader’s OD_600nm_/bioluminescence values and the value of the resident community. We defined daily invasion speed on day n (v_n_) as the distance (in wells) the invasion front moved forward on that day, and calculated the mean invasion speed v as the average daily invasion speed from day 3 until the end of the experiment or until the invader took over all wells.

### Interaction experiments

#### Overview

Precultures were prepared following a similar procedure as in invasion experiments. To get 3 biological replicates, 3 independent precultures were initiated from different colonies (for Su, Lp, and Pa) or independent aliquots from glycerol stock (for community). We initiated the interaction experiments by mixing the invader and the resident cultures at different fractions and diluting 1/200 into a 384-well Labcyte plate with 50µL IM (with corresponding buffer concentrations). The resulting interaction plate was cultured for 24h to allow species interactions. The precultures were 1/10 serially diluted and plated to obtain per volume CFU. After 24h, we measured OD_600_ (two-strain system) or bioluminescence (multi-strain system) of the interaction plate, then 1/10 serially diluted the interaction plate and plated onto corresponding plates to obtain species composition by CFU (Methods). Details are as follows.

#### Two-strain system

For each species, replicate, and buffer concentrations, precultures from day 3 were diluted 1/200X into 32 wells by transferring 250nL preculture into 50µL IM with corresponding buffer concentrations. The resulting plate was sealed with double breathable films, cultured on Mixmate at 1500rpm and 30°C for 24h, and then used to initiate interaction experiments. We started by shaking the preculture plate (1X) for 15s at 2000rpm with the Mixmate, then diluted 500nL into 49.5µL fresh IM with corresponding buffer concentrations. From the original plate (1X) and the diluted plate (1/100X), we mixed the invaders and the residents at 32 different invader ratios ranging from 0, 0.001 to 0.9, 1, and to dilute the mixture 1/200X into 384-well Labcyte plate with 50µL IM (with corresponding buffer concentrations), as detailed in Supplementary Table 4. After the transfers, we sealed the interaction plate with two layers of breathable films and incubated them in Mixmate at 1500rpm and 30°C for 24h. 8 wells of precultures from each species, replicate, and buffer concentrations were 1/10x serial diluted and plated on pH 5 (for Lp) and pH 9 (for Su) NM agar plates to obtain CFU-based initial population size.

After 24h, OD600nm and CFU-based species composition were measured for the entire interaction plate (Methods). We carried out the serial dilution for CFU, for each step, we diluted 2.5µL of culture from the previous dilution into 22.5µL PBS. The diluted plates were first mixed using Mixmate at 2000rpm for 15s, then for each well 5µL of liquid was plated onto pH 5 (for Lp) or pH 9 (for Su) NM agar plates to obtain species composition by CFU (Methods).

#### Multi-strain system

For each species/resident community and replicate, precultures from day 3 were diluted 1/200X into 32 wells with 50µL IM (10mM buffer) using Viaflo384. The resulting plate was sealed with double breathable films, cultured on Titramax 1000 shaker at 1350rpm and 30°C for 24h, and then used to initiate interaction experiments. We started by shaking the preculture plate (1X) for 15s at 2000rpm with the Mixmate, then diluted 5µL into 45µL fresh IM (10mM buffer) using Viaflo384. From the diluted plate (1/10X), we mixed the invaders and the residents at 25 different invader ratios ranging from 0.01 to 0.9 and to further dilute the mixture 1/20X (1/200X in total) into a 384-well Labcyte plate with 50µL IM (10mM buffer), as detailed in Supplementary Table 5. After the transfers, we sealed the interaction plate with two layers of breathable films and incubated them in Titramax 1000 shaker at 1350rpm and 30°C for 24h. 16 wells of precultures from each species/resident community and replicate were 1/10X serially diluted and plated on pH 7 with gentamicin (for Pa) and pH 7 without gentamycin (for community) NM agar plates to obtain CFU-based initial population or community size.

After 24h, bioluminescence and CFU-based species composition were measured for the entire interaction plate (Methods). We carried out the serial dilution for CFU with Viaflo384, for each step, we diluted 5µL of culture from the previous dilution into 45µL PBS. The serial diluted plates were mixed using Mixmate at 2000rpm for 15s, then 5µL of liquid was plated onto pH 7 with gentamicin (for Pa) or pH 7 without gentamycin (for community) NM agar plates and used for obtaining species composition by CFU (Methods).

### Obtaining interaction curves from interaction measurements

To derive interaction curves from interaction measurements, the units for initial and final invader fractions should be the same. For our experimental setup, it is thus necessary to convert between CFU-based invader fraction *f*_*CFU*_ and culture volume-based invader fraction *f*_*VOL*_, as below. The initial invader fraction was first defined by volume as the fraction of invader preculture to be mixed with the resident preculture (*f*_*VOL*_). Using measured CFU per volume data for the invader (*K*_*vd*_) and resident precultures (*K*_*rsd*_), the CFU-based invader fraction could be calculated as 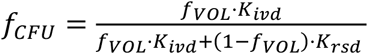. The final invader fraction was first measured based on CFU (*f*), then the volume-based *f*_*VOL*_ could be calculated by reversing the above relationship. Thus, interaction curves could be expressed either by CFU fraction or volume fraction (Supplementary Fig. 8-9) and could be converted to each other. Which of these two versions of the interaction curve to use for prediction depends on how the dispersal rate is defined. For instance, our invasion experiments defined dispersal rate as the volume of liquid transfer, so the volume-based interaction curves were used to predict invasion dynamics.

The functional form of interaction curves *g*() used in predictions were obtained by linear interpolation of experimentally measured as well as model-simulated initial and final invader fractions. The only exception was in Supplementary Fig. 15 where we inferred *g*() from a small number of data points, there we used Pchip interpolation^109^ to fit the data with a smooth and monotonic curve.

### Generalized consumer-resource model

In our generalized consumer-resource model, species interactions are mediated by biochemicals in the environment, which include both resources and toxins. Different species use, secrete, and are inhibited by different biochemicals, allowing all major types of bacterial interactions: resource competition, cross feeding, inhibition by toxins, and cross protection.

Denoting the population size of species i as *N*, the growth of species is described by:

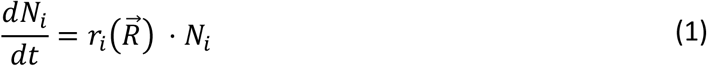

in which 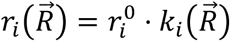 is the realized growth rate of species i, determined by intrinsic growth rate of species i 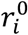 and the concentration vector of biochemicals 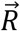 as follows:

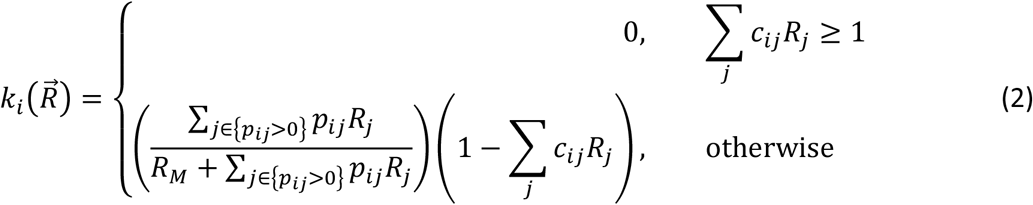

For each biochemical j, species i has a preference value *p*_*j*_ *≥* 0 and is inhibited by a coefficient

*c*_*j*_ *≥* 0. The majority of *p*_*j*_ and *c*_*j*_ are 0, meaning that species i doesn’t consume biochemical j or is inhibited by biochemical j respectively. *R*_*M*_ is the Michaelis-Menten coefficient to capture the dependency of growth rate on available biochemicals that a species can consume (i.e. resources). Motivated by experimental observation that growth stops abruptly as resources run out, a small value of *R*_*M*_ = 0.01 was used. Biochemical j can also inhibit the growth of species i and reduce its growth rate proportional to *c*_*j*_*R*_*j*_, until 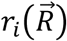 is reduced to 0.

Species can both consume and secrete biochemicals. The biochemical concentration *R*_*j*_ changes accordingly:

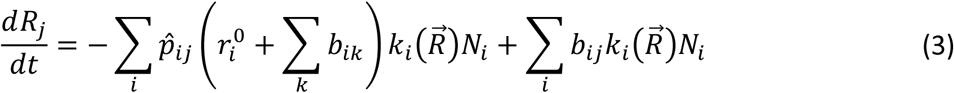

Species i secretes biochemical j at rate 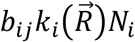 during growth. We assume an energy density of 1 for all biochemicals, i.e. we normalize biochemical concentrations by its usable energy. The normalized resource usage 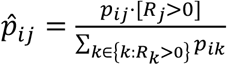, modelling a simplified scenario where species consume the available resources simultaneously and proportional to their species-specific preference to these resources. Species consume as many resources as the sum of their needs for biomass growth and secretion, so that secreting biochemicals increases resource uptake rather than reducing growth rate. This is to capture the observation that many secreted biochemicals are metabolic byproducts and would not incur a significant cost to bacteria ^110^.

### Generalized Lotka-Volterra model

In our generalized Lotka-Volterra model, each species j directly inhibits or facilitates species i, through negative and positive interaction coefficient *A*_*j*_ respectively. The growth of species i is described by

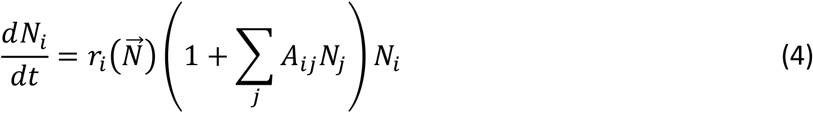

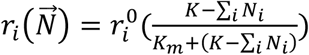 is the realized growth rate of species i, where 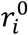 is the intrinsic growth rate of species i, *K* = 3 represents a relatively large carrying capacity, and *K*_*m*_ = 0.1 is a relatively small coefficient that sharply reduces growth rate when the community approaches carrying capacity. In this setting, 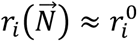 in most cases, growth is constrained by a carrying capacity shared by the community, preventing unbounded growth caused by positive interactions.

### Invasion, interaction curve, and prediction in mechanistic models

#### Random generation of resident communities and invaders

For both the generalized consumer-resource model (CR model) and the generalized Lotka-Volterra model (LV model), we generated 100 random resident communities (each starting with 50 species) and 20 random invaders. We studied the invasion dynamics for each of the 100*20 pairs. See Supplementary Text 1 for the choice and distribution of parameters.

We simulated resident communities from initial populations of 10^−3^ per species for 30 dilution cycles when most community compositions reached equilibrium. In each dilution-growth cycle, the species (and biochemicals for the CR model) were diluted by 1/200X from the previous cycle as in the experiment, for the CR model another 199/200X of supplied resources were added (1/3 a.u. for 3 biochemicals out of 30). From this new initial condition, we simulated growth and interaction as described by the mechanistic model for 24 time units, i.e. a digital day. We discarded species with population size < 10^−5^ at the end of the 30^th^ cycle and simulated 10 more daily dilution cycles to obtain the equilibrium resident communities. Similarly, we simulated invaders from an initial population of 10^−3^ for 10 dilution cycles to get the equilibrium invader populations.

#### Simulating invasion dynamics from mechanistic models

We simulated invasion dynamics from both mechanistic models similarly as in the invasion experiment. 10 wells were initiated with the equilibrium invader population, followed by 10 wells with the equilibrium resident community. We simulated 20 dispersal-dilution-growth cycles. At the beginning of each cycle n+1 (t=0), we calculated the species composition 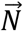 (and resource composition 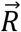 for CR model) for each well position x after dispersal with rate m and dilution with factor δ = 1/200 as:

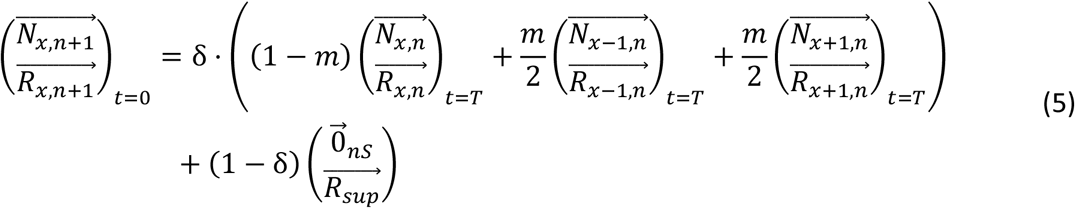

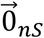 is a zero vector with a size equal to the species number and 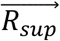 is a vector of supplied biochemical concentrations in fresh media. We simulated growth and interaction as described by the mechanistic models for T=24 time units, and used the resulting species and resource compositions (t=T) as the starting point to simulate the next dispersal-dilution-growth cycle.

#### Simulating interaction measurements

Similar to experimental setup, we simulated the interaction measurements by mixing equilibrium invader population and resident community (for CR model also biochemicals) at 29 different fractions ([0.0, 0.0001, 0.001, 0.004, 0.01, 0.02, 0.03, 0.05, 0.1, 0.15, 0.2, 0.25, 0.3, 0.4, 0.5, 0.6, 0.7, 0.75, 0.8, 0.85, 0.9, 0.95, 0.97, 0.98, 0.99, 0.996, 0.999, 0.9999, 1.0], symmetric around 0.5). Then from these different initial conditions, we simulated the mechanistic model for 1 dilution-growth cycle, specifically, we multiplied the population size (and biochemical concentrations) by δ and simulated growth and interaction as described by the mechanistic models for T=24 time units. The results were final species compositions for each invader-resident pair and different initial invader fractions.

#### Predicting simulated invasion from simulated interaction measurements

From the initial and final compositions from the simulated interaction measurements described above, we calculated the “volume”-based initial and final fraction of invaders, similar as for experimental data. We obtained the interaction curve *g*() by linearly interpolating the initial and final fraction. All details from the mechanistic model were lost in this process, and we only used the resulting 1-variable interaction curve to predict simulated invasion dynamics following the parameter-free framework in Fig. 3.

#### Numerical solver

We run all simulations in Python version 3.11 with ‘solve_ivp’ in scipy version 1.13.0. We used Runge–Kutta–Fehlberg method (RK45) with adaptive stepsize (relTol=1e-3 and absTol=1e-6).

## Supplementary Information

Supplementary Figures 1-19

Supplementary Tables 1-5

Supplementary Text 1

