## Supplementary Data and Information for "Biotic resistance predictably shifts microbial invasion regimes"

### Supplementary Figures

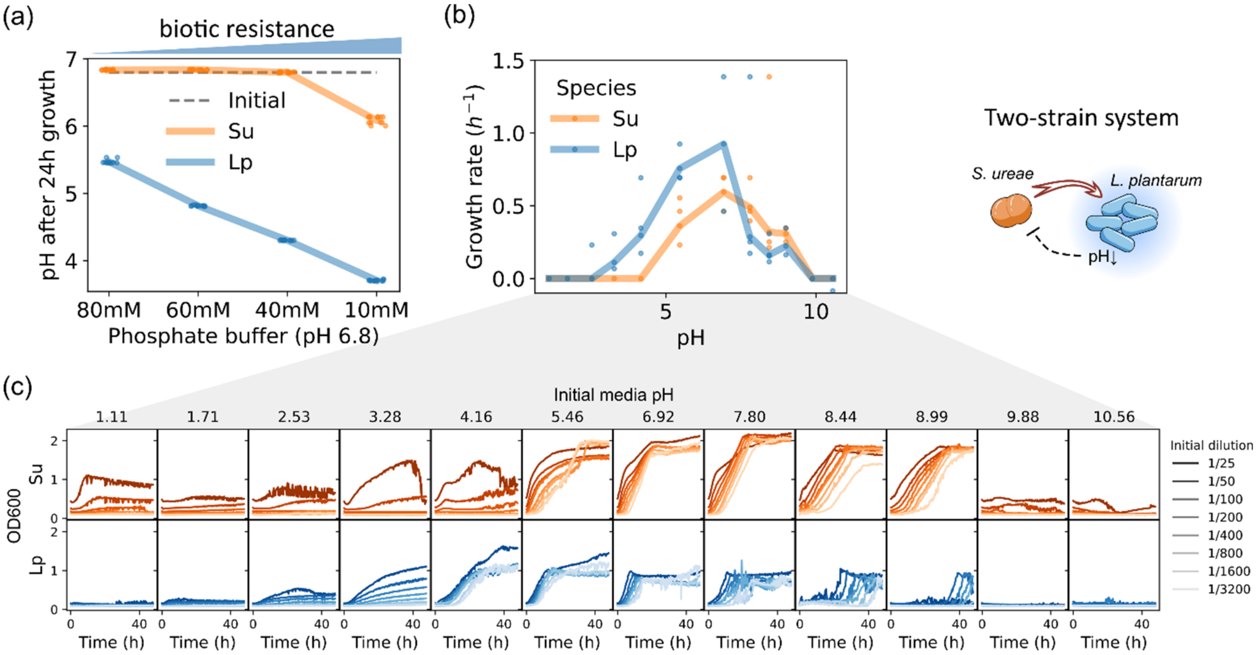

**Supplementary Figure 1. Building the two-strain system with tunable biotic resistance mediated by pH. (a)** pH of invader (Su) and resident (Lp) bacteria cultures after 24h of growth (N=12) in IM media with different buffer concentrations. A decrease in buffer concentration causes an increased pH shift by the growing bacteria, notably pH reduction by Lp. **(b)** Exponential growth rates of Su and Lp growth at different initial pH values, a lower pH inhibits Su more than Lp. Values were calculated from growth curve measurements in (c), lines show mean values (N=1-4, see below). **(c)** Growth curves for Su (orange) and Lp (blue) in IM (80mM buffer) at different initial pH values, measured by $OD_{600}$. Bacteria precultures were 1/2X serially diluted from saturated precultures to inoculate growth curve measurements with different initial population sizes (initial dilution, dark to light). The exponential growth rate in (b) was calculated as $r = \frac{\ln(2)}{t_{double}}$, where doubling time $t_{double}$ was measured as the time difference for a pair of adjacent growth curves to reach a threshold of 0.5 (for Su) or 0.2 (for Lp). Each growth curve was only involved in one calculation pair to ensure independence. Depending on how many growth curves cross the threshold, 1-4 $r$ values were obtained for each pH.

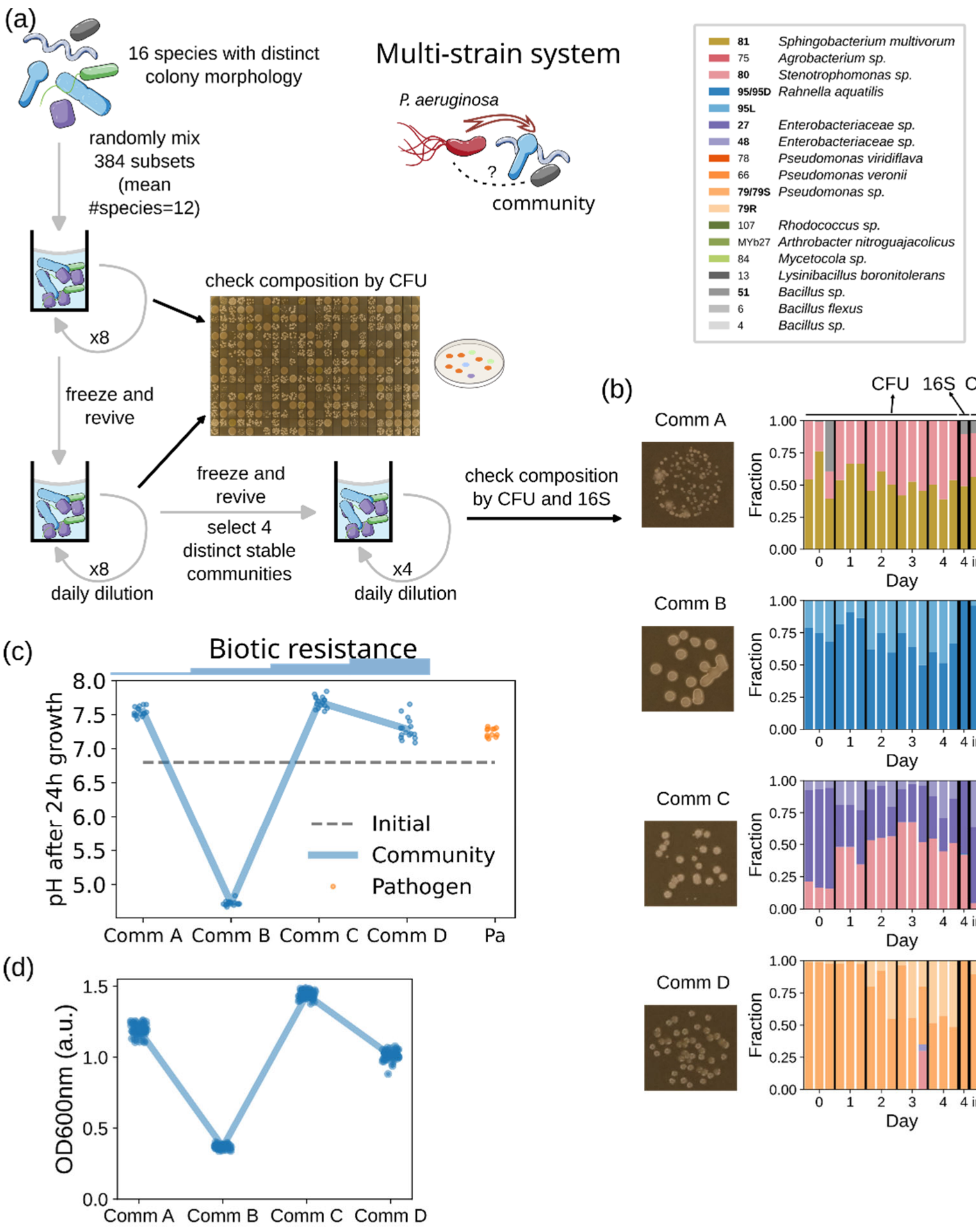

**Supplementary Figure 2. Building the multi-strain system with an invading pathogen and randomly assembled microbial communities. (a)** Building stable resident communities from *C. elegans* gut strains (Methods). Strains that remain detectable in obtained communities are highlighted in bold. **(b)** Resident community compositions over 4 days of 1/200X daily dilution after reviving from glycerol stock (days 0-4), for 3 replicates each initiated from an independent inoculum. For each resident community, compositions of 3 wells on day 8 of the invasion experiment (inv8) are also shown. The 3 wells from invasion experiment were taken from the 3 lowest dispersal rates and the farthest from the invasion front, thus least influenced by invader – they only encountered the invader for 1 day. Species compositions were obtained by counting colonies with distinct morphologies (CFU) and additionally by 16S sequencing on day 4.

Upon culturing, colonies of strain 95 diverged into dark (95D) and light (95L) morphologies, which were not separable by streaking out single colonies, i.e. non-heritable heterogeneity; colonies of strain 79 diverged into smooth-flat (79S) and rough-bumpy (79R) morphologies, streaking out single colonies gave rise to colonies only of the same morphology, i.e. heritable heterogeneity. **(c)** pH of resident community culture after 24h doesn't show a correlation with biotic resistance, contrary to the simple system (Supplementary Fig.1a). **(d)** Carrying capacity (OD600nm) of resident community culture on day 1 of invasion didn't show a correlation with biotic resistance.

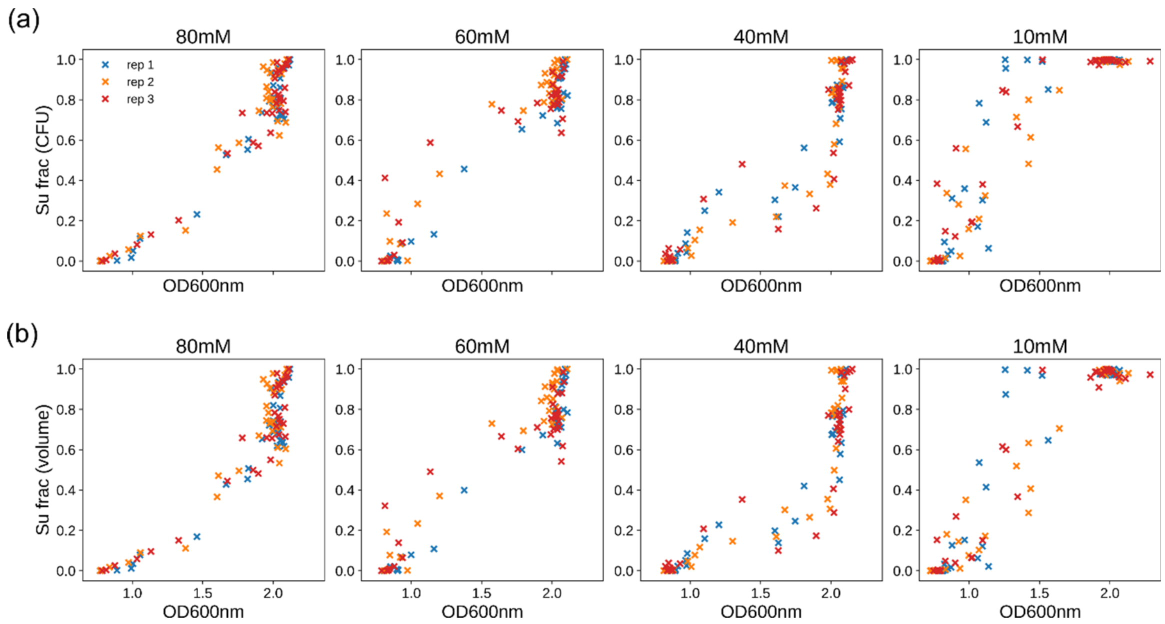

**Supplementary Figure 3. In the two-strain system invasion can be tracked by measuring $OD_{600}$.** Invader (Su) fraction in terms of CFU (a) and equivalent volume of saturated invader and resident species cultures (Methods) (b) both correlate with OD600nm measurement, for the four phosphate buffer concentrations used. Rep1-3 are three biological replicates from different single colonies.

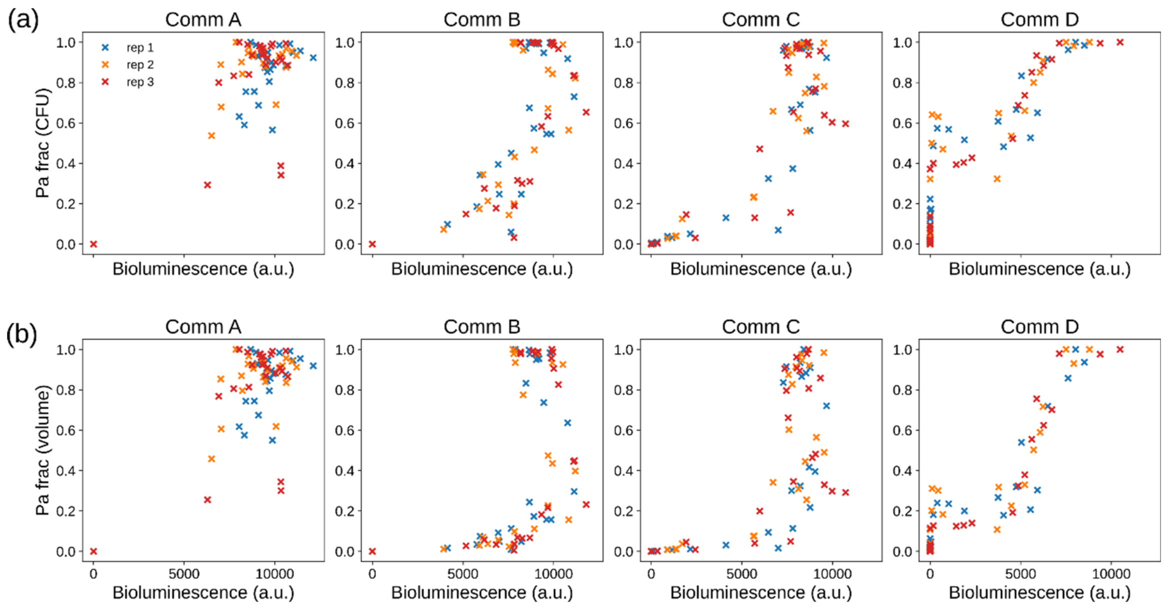

**Supplementary Figure 4. In the multi-strain system invasion can be tracked by measuring bioluminescence.** Invader fraction (Pa) in terms of CFU (a) and equivalent volume of saturated invader and resident cultures (Methods) (b) both correlate with bioluminescence measurement, for invasion into the four resident communities used. The pathogen and invader Pa was tagged with luciferase to produce bioluminescence. Rep1-3 are three biological replicates from different single colonies of Pa and different inoculum of resident communities.

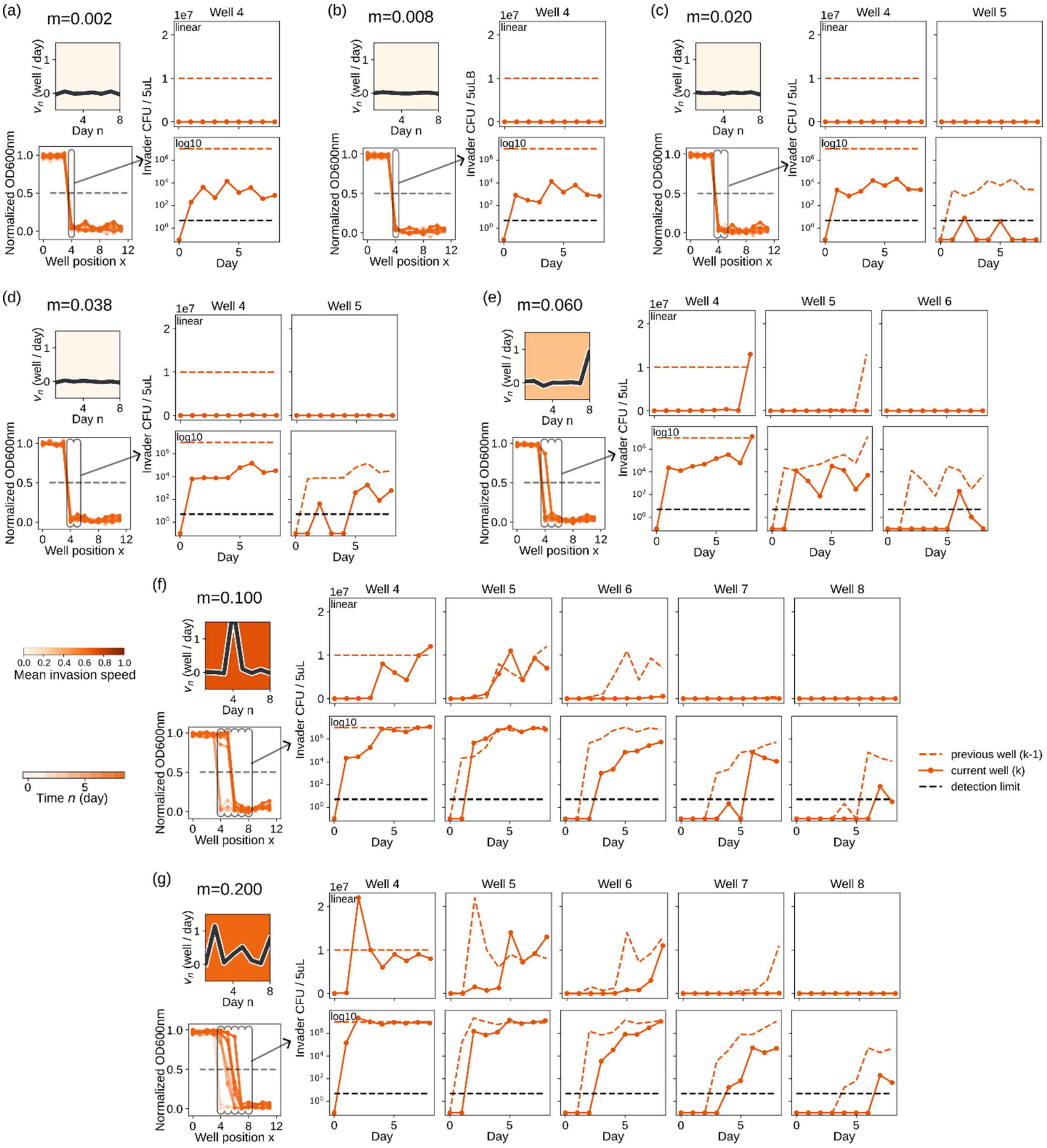

**Supplementary Figure 5. Dynamics of the absolute invader (Su) population in pinned and pulsed invasions.** Each panel shows results from a different dispersal rate for the two-strain system with 40mM phosphate buffer (intermediate resistance). **(a-d)** In pinned invasion, the invader was permanently present in the invaded wells but unable to fully establish. Upper left: daily and mean invasion speeds; lower left: invasion waves; both are as described in Main Fig. 2b. Right: daily absolute invader population size measured by CFU, presented in both linear (upper) and log10 (lower) scales, from the first well being invaded (well 4) to the last well with a detectable invader population. **(e-g)** In pulsed invasion, the invader slowly accumulated at the invasion front before suddenly taking over. Subpanels are described as above.

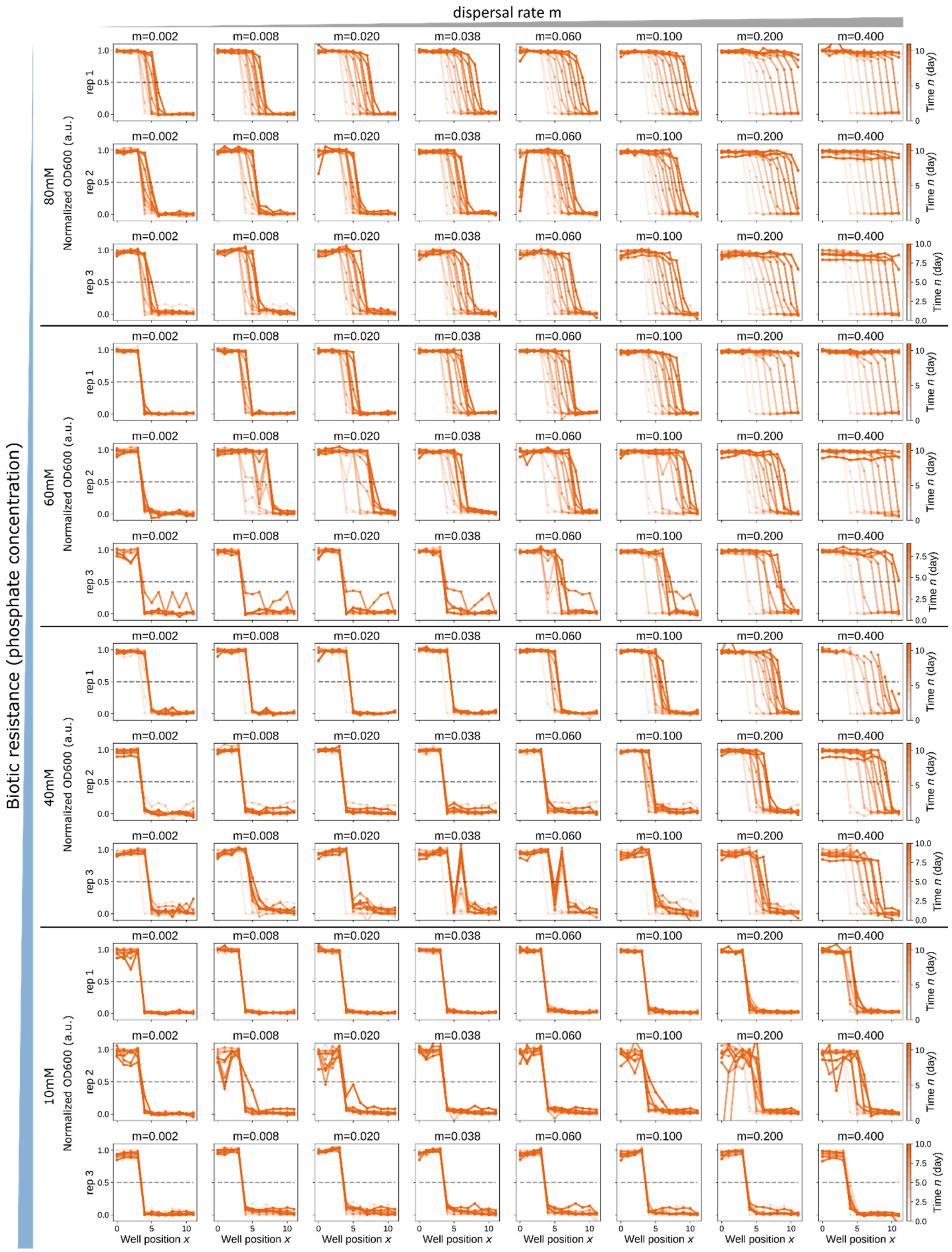

dispersal rate m
Biotic resistance (phosphate concentration)
80mM
60mM
40mM
10mM
Normalized OD600 (a.u.)
rep 1
rep 2
rep 3
m=0.002
m=0.008
m=0.020
m=0.038
m=0.060
m=0.100
m=0.200
m=0.400
Time n (day)
Well position x

**Supplementary Figure 6. Raw invasion waves for the two-strain system.** Invasion waves as described in main Fig. 2 for 4 biotic resistance levels resulting from different phosphate buffer concentrations. 3 biological replicates for each resistance level, each with 8 dispersal rates.

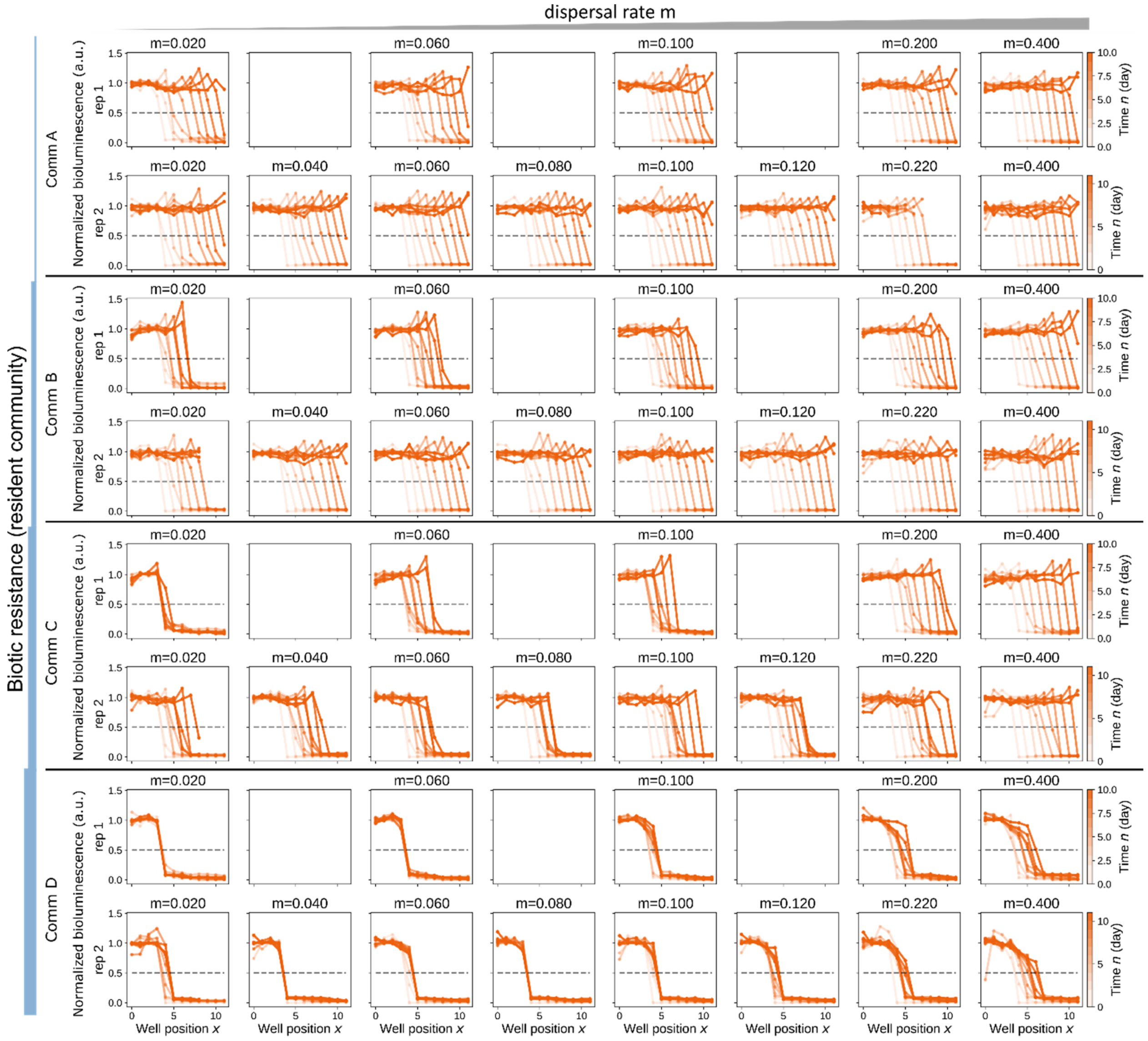

**Supplementary Figure 7. Raw invasion waves for the multi-strain system.** Invasion waves as described in main Fig. 2, for 4 resident communities with different resistance levels. 2 biological replicates for each resistance level, one with 5 and another with 8 dispersal rates, due to technical failure during the experiment for replicate one.

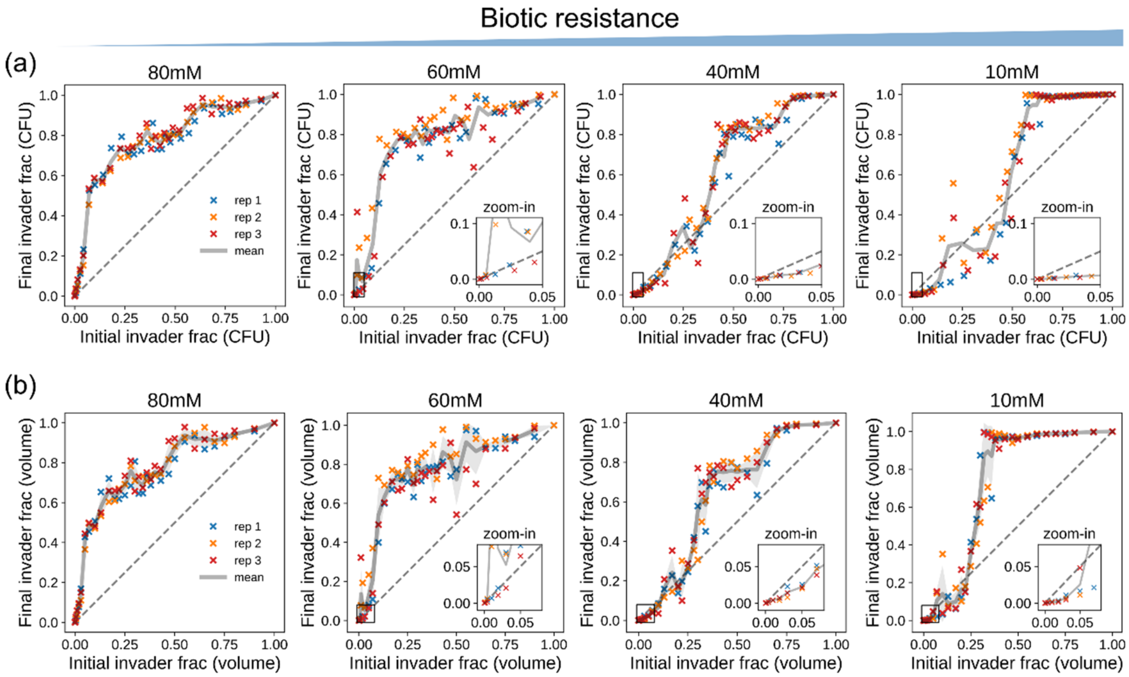

**Supplementary Figure 8. Interaction curves for the two-strain system.** Relationship between the initial and final invader fraction after 24h culture, as described in main Fig. 3a. **(a)** Invader (Su) fraction as represented by fraction of CFU, which is also shown in main Fig. 3b. **(b)** Invader (Su) fraction as represented by fraction of volume when mixing bacteria cultures (Methods). As our experimental setup defined dispersal rate in terms of volume instead of CFU, the predictions were made from the interaction curves in (b). The grey line shows the averaged interaction curve, light grey shadow shows curves of mean interaction ±1 standard deviation.

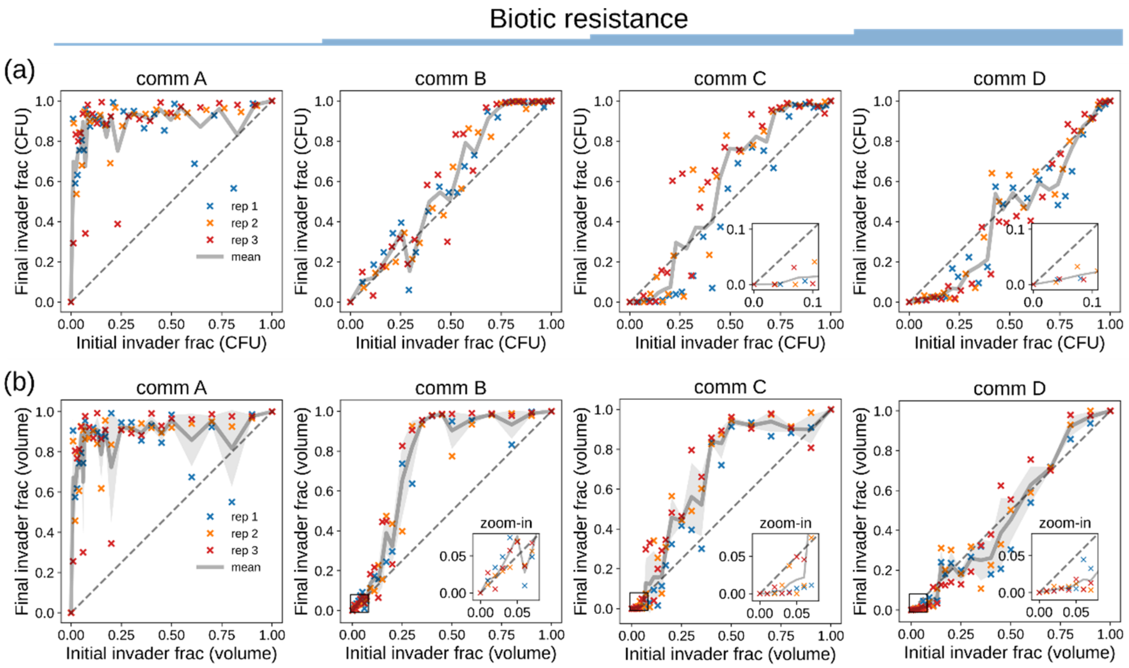

**Supplementary Figure 9. Interaction curves for the multi-strain system.** Relationship between the initial and final invader fraction after 24h culture, as described in main Fig. 3a. **(a)** Invader (Pa) fraction as represented by fraction of CFU, which is also shown in main Fig. 3c. **(b)** Invader (Pa) fraction as represented by fraction of volume when mixing bacteria cultures (Methods). As our experimental setup defined dispersal rate in terms of volume instead of CFU, the predictions were made from the interaction curves in (b). The grey line shows the averaged interaction curve, light grey shadow shows curves of mean interaction ±1 standard deviation.

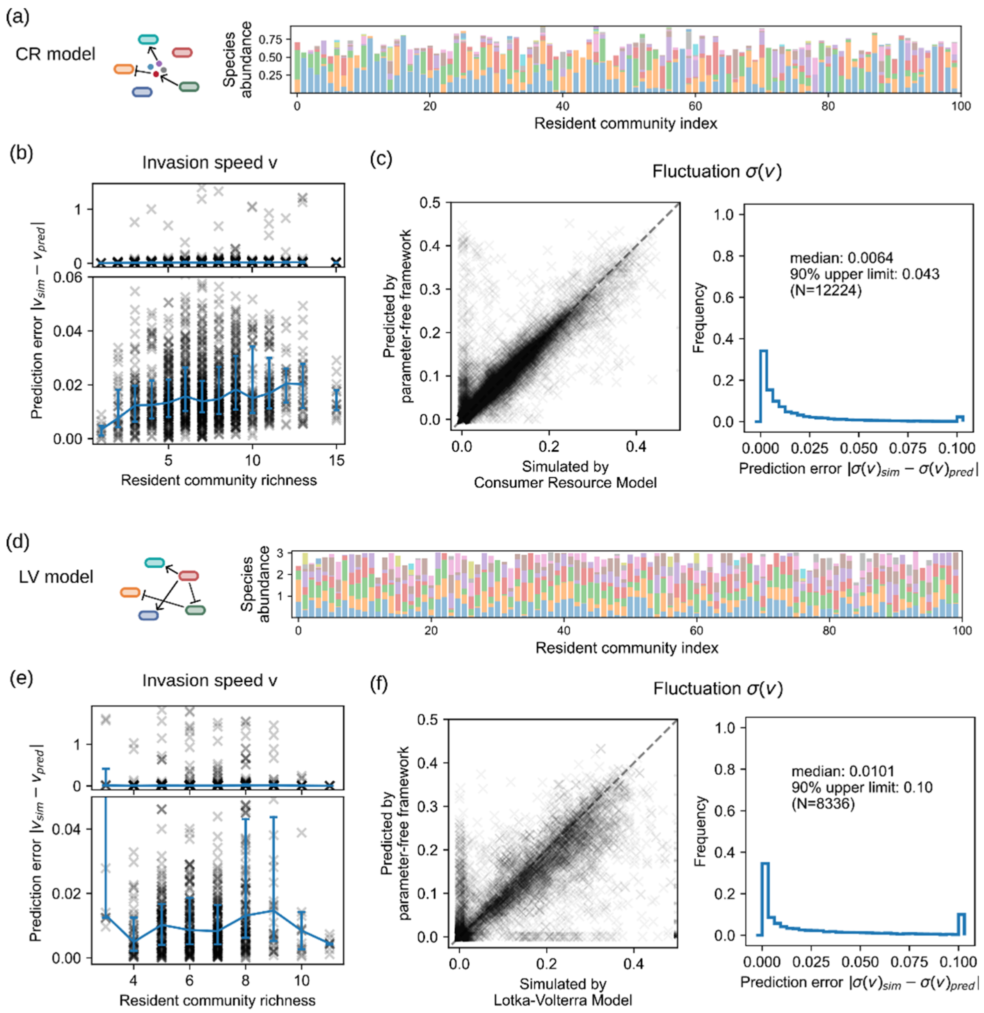

**Supplementary Figure 10. Parameter-free framework accurately predicts invasion dynamics simulated by mechanistic models. (a, d)** Resident community compositions for generalized consumer-resource model (a) and generalized Lotka-Volterra model (d). The same color in different resident communities generally represents different species. **(b, e)** Prediction error for mean invasion speed v remains low regardless of the richness of resident communities for both models. The median (blue line) and quantiles (blue bars) are shown. **(c, f)** Fluctuation of invasion speed, defined as the standard deviation of daily invasion speed $\sigma(v)$, is correctly predicted for most cases with non-negative invasion speed for both models.

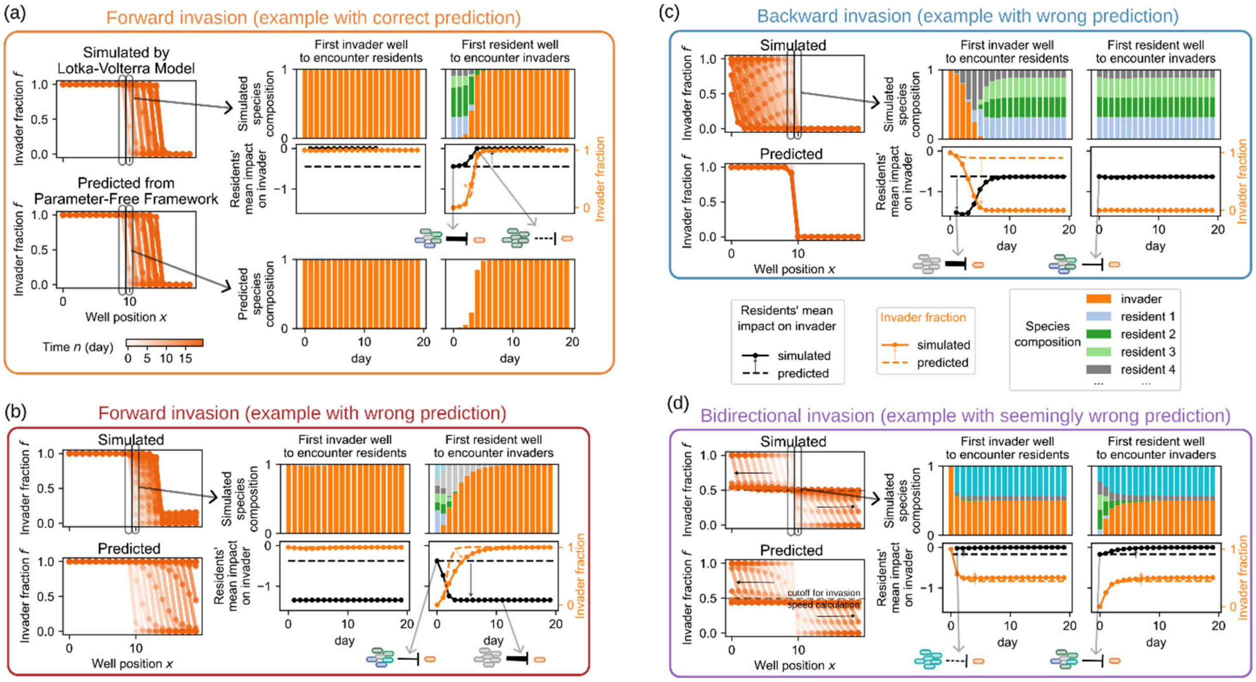

**Supplementary Figure 11. Examples of simulated invasion demonstrate the applicability and limitation of the parameter-free framework.** Four examples are taken from the generalized Lotka-Volterra Model, also highlighted in Supplementary Fig. 12b. For each example, invasion waves (dispersal rate m=0.06) from both simulation and prediction are shown on the left. For the two wells that were initially located on the border of the invader and the resident community, daily species composition and the disturbed resident community's mean impact on invader $\alpha_{0\cdot} = \frac{\sum_j \alpha_{0j} \cdot N_j}{\sum_j N_j}$ are shown on the right. $\alpha_{0j}$ is the interaction coefficient of resident species j on invader in the generalized Lotka-Volterra Model; positive and negative values indicate facilitation and inhibition, respectively. $N_j$ is the population size of resident species j. **(a)** An example of forward invasion with correct prediction (resident #0, invader #4). Invasion slightly reduces residents' inhibition towards the invader (solid black line), which facilitates invasion only minorly (solid orange line and orange arrow) because the invader already reaches a high fraction; thus, prediction (dashed orange line) remains accurate. **(b)** A rare example of forward invasion with wrong prediction (resident #51, invader #18). Invasion significantly increases the resident's inhibition towards the invader (solid black line) while invader fraction is still low, making the simulated invasion (solid orange line) slower than predicted (dashed orange line). **(c)** An example of backward invasion (resident #22, invader #4) with wrong prediction. The small fraction of resident species that initially invades back into the invader population is of distinct composition and shows much stronger inhibition towards the invader (solid black line). Consequently, while the invasion dynamics are predicted to be pinned, the resident community intrudes backward in simulation. **(d)** An example of bidirectional invasion (resident #87, invader #18). The invader coexists with some of the resident species, so while the invader moves forward, part of the resident community also travels backward. In such scenarios, invasion speed depends on the invader fraction threshold used for calculation, and as an artifact small discrepancy between predicted and simulated dynamics leads to a seemingly large difference in predicted vs. simulated invasion speed (Supplementary Fig. 12b, vii).

(a)

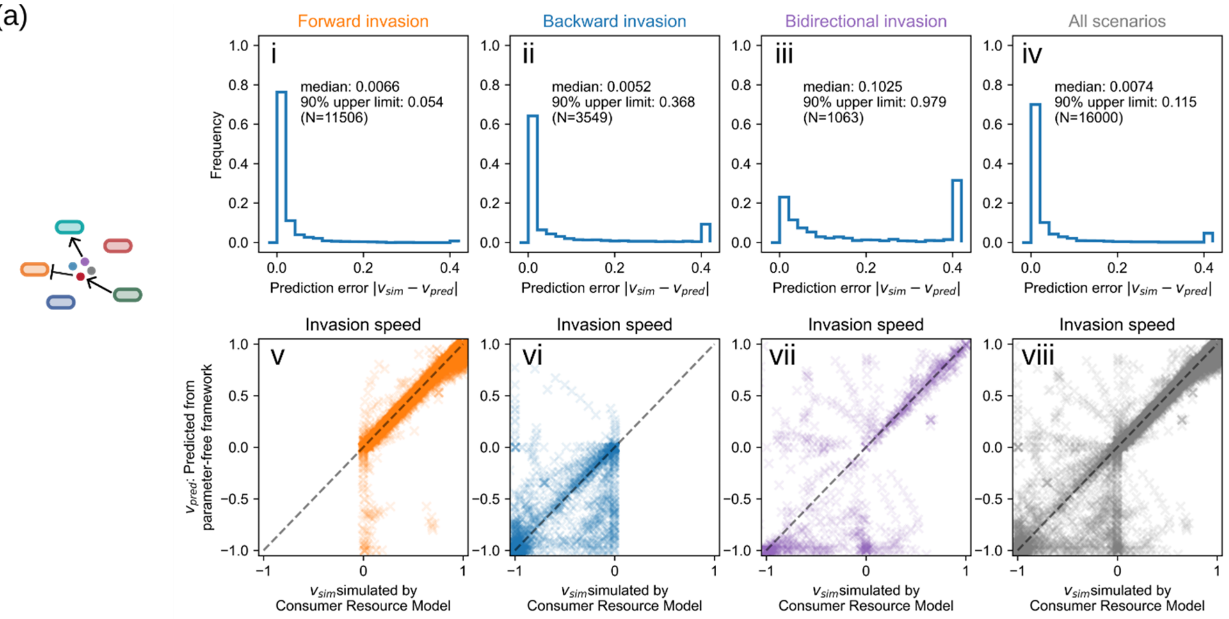

(b)

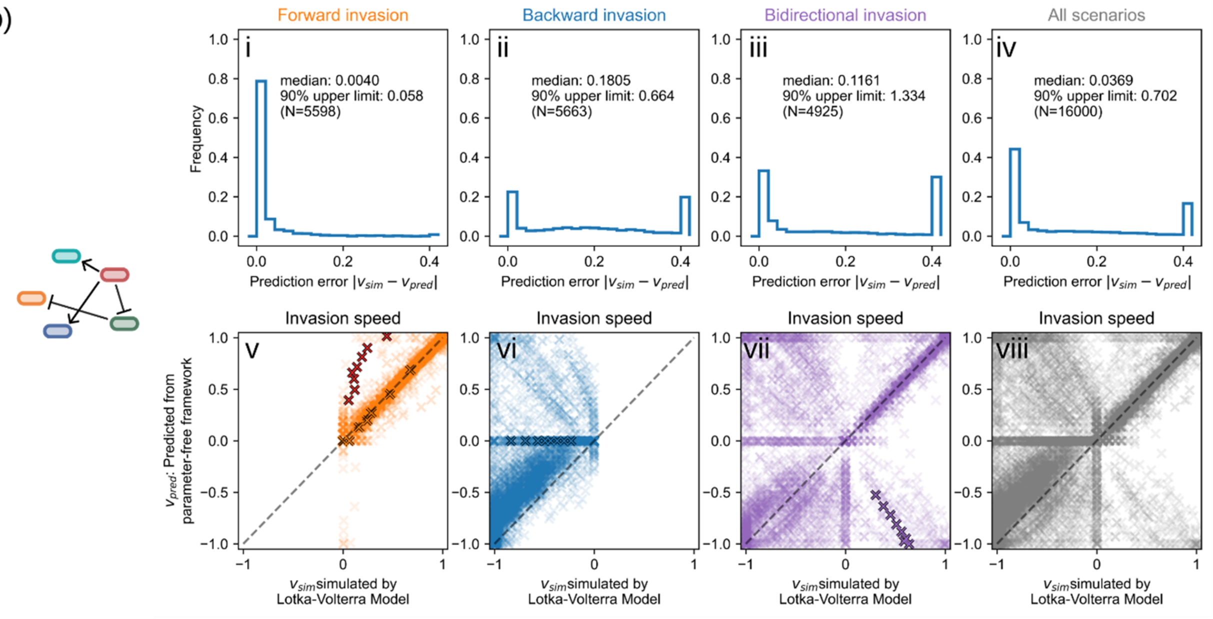

**Supplementary Figure 12. Prediction is highly accurate for forward invasion but less for backward and bidirectional invasions in mechanistic models.** Predicted vs. simulated invasion speed (v-viii) and the distributions of prediction errors (i-iv) for generalized consumer resource model (a) and Lotka-Volterra model (b). Values for all 8 dispersal rates from the examined examples in Supplementary Fig. 11 are marked by black-outlined crosses in (b) v-vii. Predictions for forward, backward, and bidirectional invasion (as defined in Supplementary Figure 11) are shown separately. For forward invasions, the prediction is very accurate (i, v, orange). For backward invasion the predictions quite often deviate quantitatively from simulations (ii, vi, blue), but mostly still predict invasion to be backward or pinned. This is somewhat expected, as during backward invasion, the resident community frequently shifts to a different composition at the backward invasion front, as demonstrated in Supplementary Fig. 11c. For bidirectional invasion, the predicted invasion speed is generally not trustable (iii, vii, purple), which is mostly due to an artifact originated from the definition of invasion speed, as demonstrated in Supplementary Fig. 11d. The predictions mostly capture the qualitative bidirectional waves and their speed for the forward part, but the speed for the backward part as well as the coexistence threshold is frequently not accurately predicted.

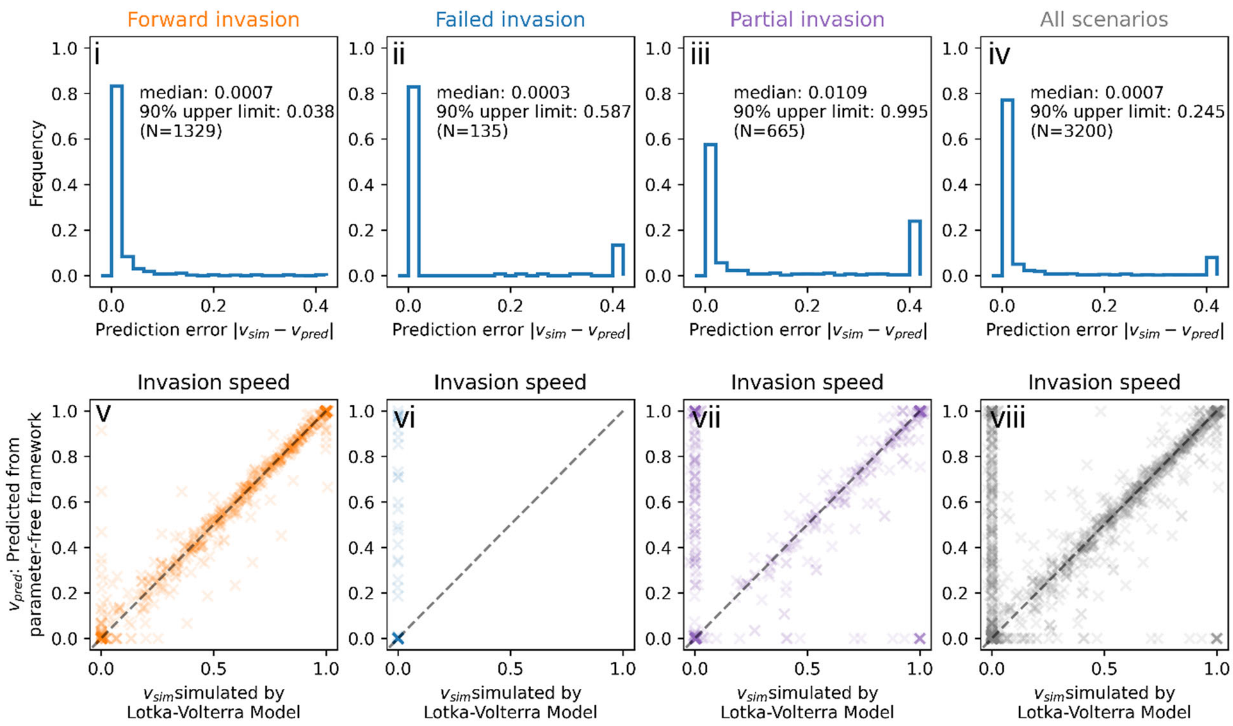

**Supplementary Figure 13. Prediction is highly accurate also when dispersal is unidirectional.** Predicted vs. simulated invasion speed (v-viii) and the distributions of prediction errors (i-iv) for generalized Lotka-Volterra model and unidirectional dispersal. In addition, predictions for the three distinct scenarios are shown separately: (1) forward invasion (i, v, orange), where invader takes over and the invasion front moves forward; (2) failed invasion (ii, vi, blue), where the invader could not grow within the resident community and the invasion front does not move; and (3) partial invasion (iii, vii, purple), where the invader coexists with the resident community, since the invader cannot exclude the residents, only part of the invasion front moves forward. These scenarios are analogous to forward, backward including pinned, and bidirectional invasions when dispersal is bidirectional as shown in Supplementary Figs. 11-12.

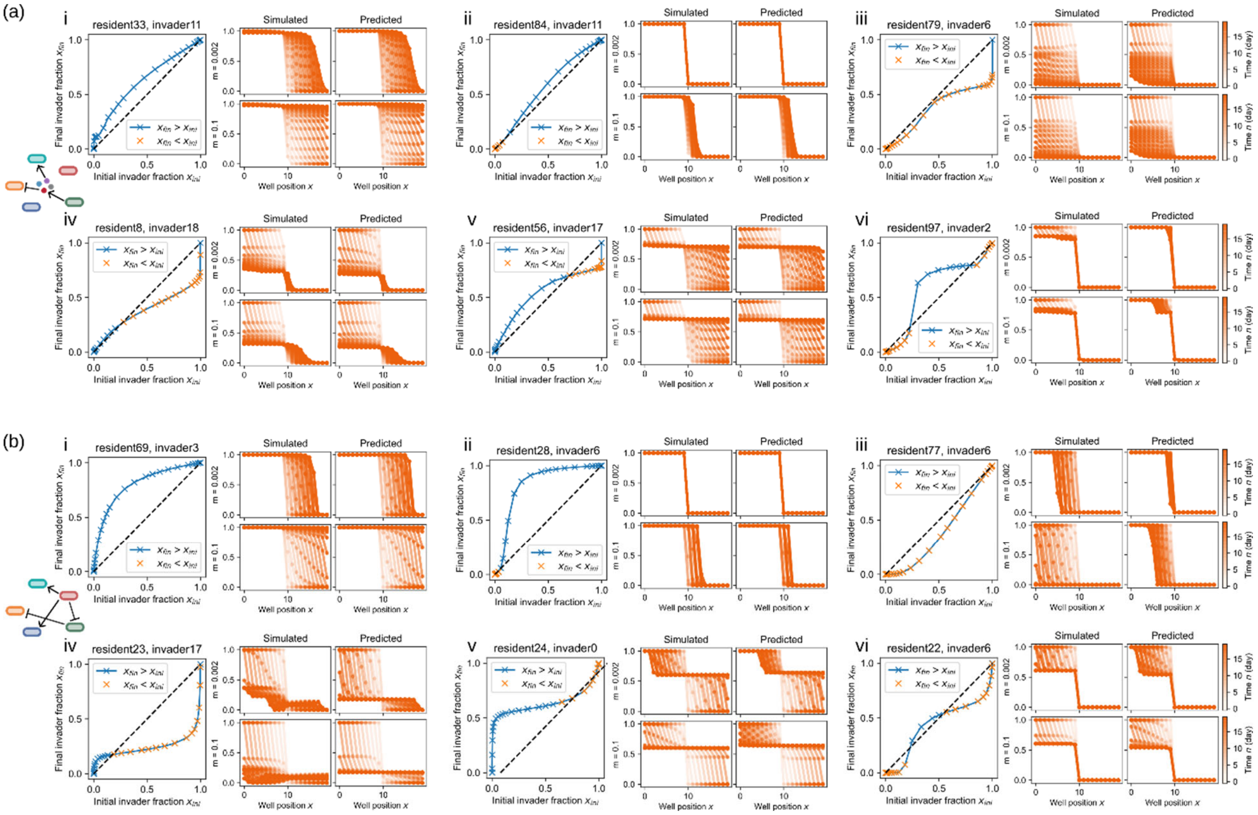

**Supplementary Figure 14. Example interaction curves and invasion dynamics from mechanistic models. (a)** Examples from generalized consumer-resource model. **(b)** Examples from generalized Lotka-Volterra model. Each subpanel shows on the left the interaction curve between a given invader and resident community, and on the right, simulated and predicted invasion dynamics for a low (m=0.002) and a high (m=0.1) dispersal rate. **i**: interaction curve consistently above diagonal results in forward invasion regardless of dispersal rate. **ii**: interaction curve that is below diagonal for low initial invader fraction $x_{ini}$ and crosses above the diagonal for high $x_{ini}$ results in pinned invasion at a low dispersal rate and forward invasion at a high dispersal rate. A critical dispersal rate emerges in this scenario. **iii**: interaction curve consistently below diagonal results in backward invasion regardless of dispersal rate. **iv-vi**: interaction curve that is above diagonal for low initial invader fraction $x_{ini}$ and crosses below the diagonal for high $x_{ini}$ results in the coexistence of invader and resident species and, consequently, bidirectional invasion. The coexistence fraction between the invader and the resident in bidirectional invasion depends on the position of crossing.

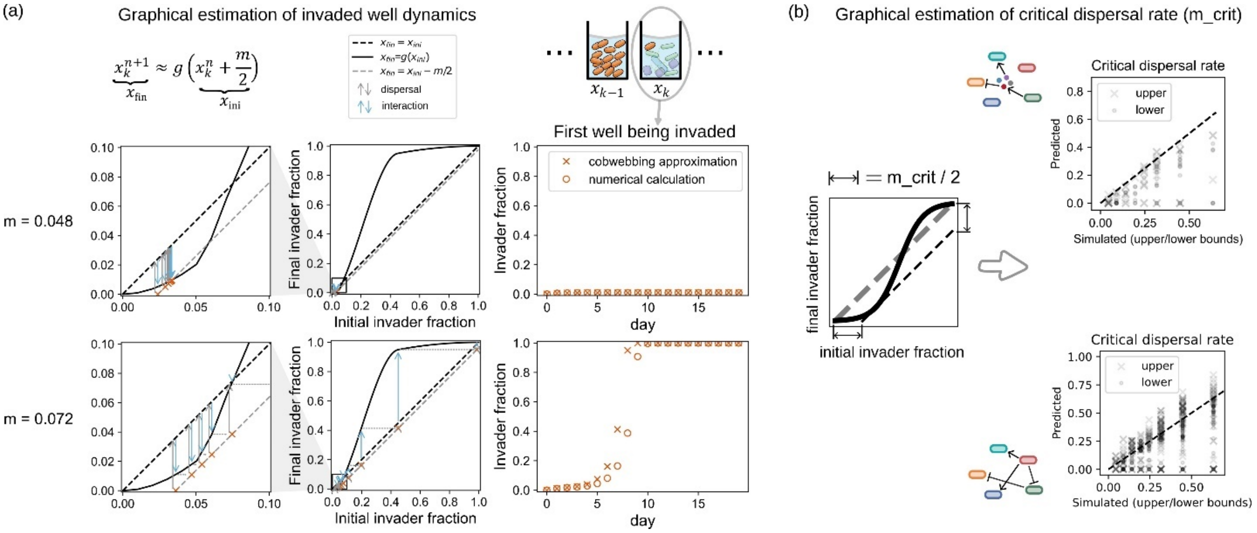

**Supplementary Figure 15. The parameter-free framework can be graphically approximated using cobwebbing. (a)** Graphical estimation of the invader population dynamics in the first well being invaded, for dispersal rates smaller (upper panels) and bigger (lower panels) than the critical dispersal rate $m_{crit}$. Assume well k is the first resident well to be invaded. The invader dynamics of well k ($x_k^n$) as shown in Main Fig. 3a can be approximated by $x_k^{n+1} \approx g\left(x_k^n + \frac{m}{2}\right)$ under the assumptions that dispersal rate is small ($m \ll 1$), the invader population is small in invaded wells ($x_k^n, x_{k+1}^n \ll 1$), and the resident populations in the invader population are small ($1 - x_{k-1}^n \ll 1$). With these approximations the dynamics of $x_k^n$ could be visualized by a modified version of cobwebbing (left panels), in which grey arrows from shifted diagonal $x_{fin} = x_{ini} - \frac{m}{2}$ describe the effects of dispersal, and blue arrows towards the interaction curve $x_{fin} = g(x_{ini})$ describe the effects of biotic resistance/species interactions. The dynamics of invader fraction in invaded well $x_n^k$ obtained by this graphical approach is similar to the numerical calculation made by using the original equation in Main Fig. 3a, especially during the initial phase of invasion when $x_n^k$ is still small (right panels). **(b)** The graphical approach in (a) indicates that the critical dispersal rate $m_{crit}$ is two times the maximum reduction of invader fraction ($x_{ini} - x_{fin}$) for the lower part of the interaction curve (left panel). This prediction is confirmed by comparing graphically obtained $m_{crit}$ to simulated values in both generalized consumer-resource model (upper right panel) and generalized Lotka-Volterra model (lower right panel). Because the invasions are simulated under 8 discrete dispersal rates, only upper and lower limits of $m_{crit}$ can be obtained. Taken together, while the graphical approach here only holds true under limited conditions, it provides important intuition about how the shape of the interaction curve impacts invasion dynamics.

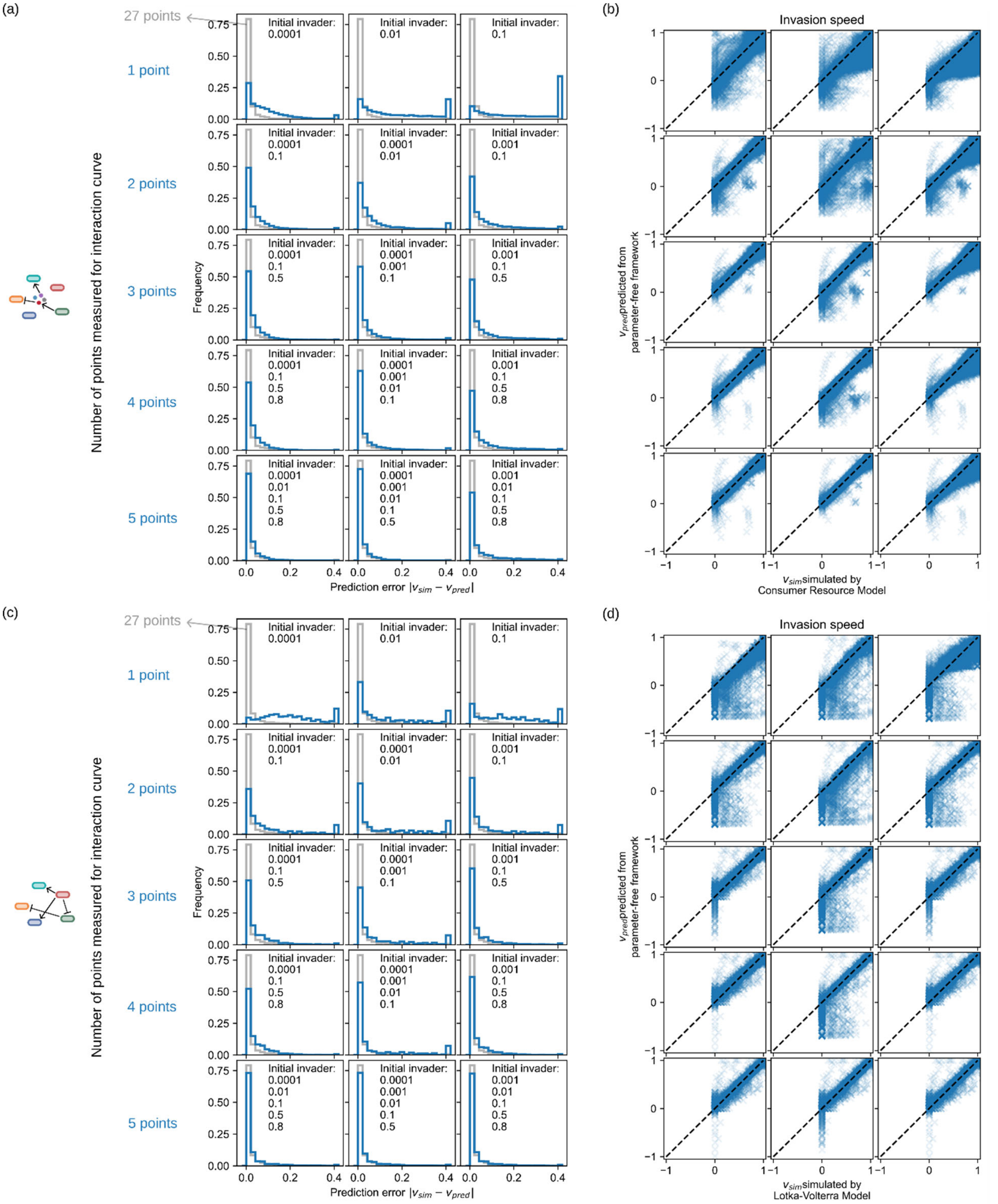

**Supplementary Figure 16. A few points are enough to approximate interaction curves and give accurate predictions for simulated invasions.** Prediction accuracy of invasion speed when the interaction curves are approximated from only 1-5 points. **(a-b)** Prediction error of invasion speed (a) and predicted vs. simulated invasion speed (b)

for generalized consumer-resource model. Blue text labels the number of points (1-5) used to approximate interaction curves for each row. For each given number, results from three sets of different points are shown, as annotated by initial invader fraction in each subpanel. For example, “initial invader: 0.001, 0.1” means only final invader fraction data $x_{fin}$ for initial invader fractions $x_{ini} \in \{0.001, 0.1\}$ are used to approximate the interaction curves, instead of all 27 points as shown in main text and also in grey here. The approximation is done by Pchip interpolation(Fritsch and Butland, 1984) of $(x_{ini}, x_{fin})$ from the chosen points plus the two ends of the interaction curves (0, 0) and (1, 1). **(c-d)** Similar to (a-b) but for generalized Lotka-Volterra model. For both models the results also demonstrate that the prediction would generally be more accurate if the initial invader fractions used cover a relatively larger range of values (the left and right columns in comparison to the middle column), supporting the idea that the overall shape of the interaction curve matters for invasion.

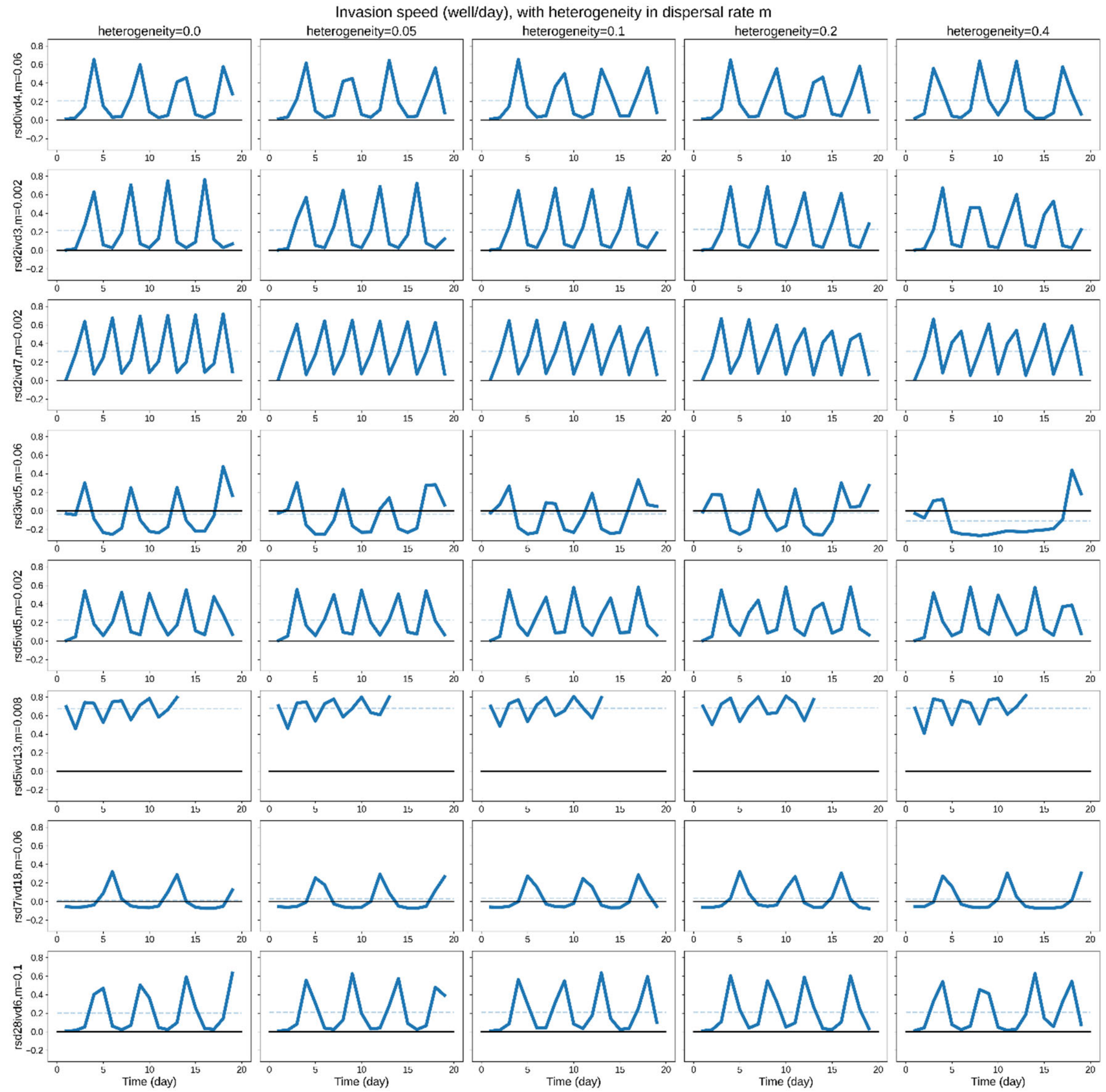

**Supplementary Figure 17. Daily invasion speed for invasion along patches with heterogeneous dispersal rate m.** Each row shows daily invasion speed (solid blue line) and mean invasion speed (horizontal blue dash) for an example with the given resident community (rsd), invader (ivd) and dispersal rate (m). The heterogeneity of dispersal rate m across patches increases from left to right. For given heterogeneity level H, the dispersal rate m out of each patch is multiplied by a random value independently drawn from the uniform distribution [1-H, 1+H], whereas H=0 means no heterogeneity. To compare better with previous theoretical results, we defined the front position as the sum of invader fractions (divided by carrying capacity 1) when computing the invasion speed instead of the position above the 50% threshold; the conclusions stayed the same regardless.

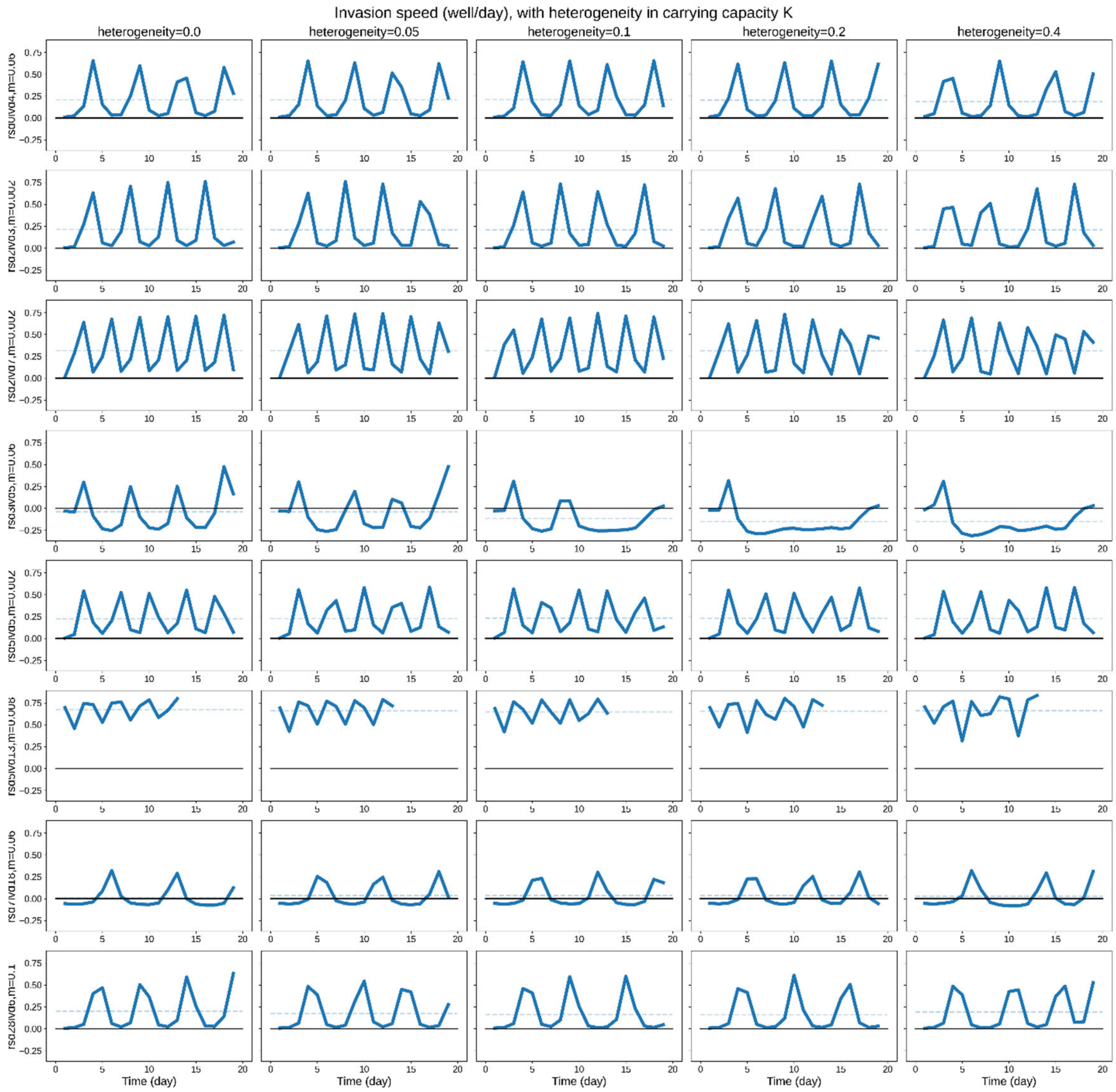

**Supplementary Figure 18. Daily invasion speed for invasion along patches with heterogeneous carrying capacity K.** Each row shows daily invasion speed (solid blue line) and mean invasion speed (horizontal blue dash) for an example with the given resident community (rsd), invader (ivd) and dispersal rate (m). The heterogeneity of carrying capacity K across patches increases from left to right. For given heterogeneity level H, the carrying capacity of each patch is independently drawn from the uniform distribution [1-H, 1+H], H=0 means no heterogeneity. Since we normalized carrying capacity in Lotka-Volterra model to 1, this is equivalents to divide the interaction terms \alpha by these randomly drawn K. To compare better with previous theoretical results, we defined the front position as the sum of invader fractions (divided by carrying capacity 1) when computing the invasion speed instead of the position above the 50% threshold; the conclusions stayed the same regardless.

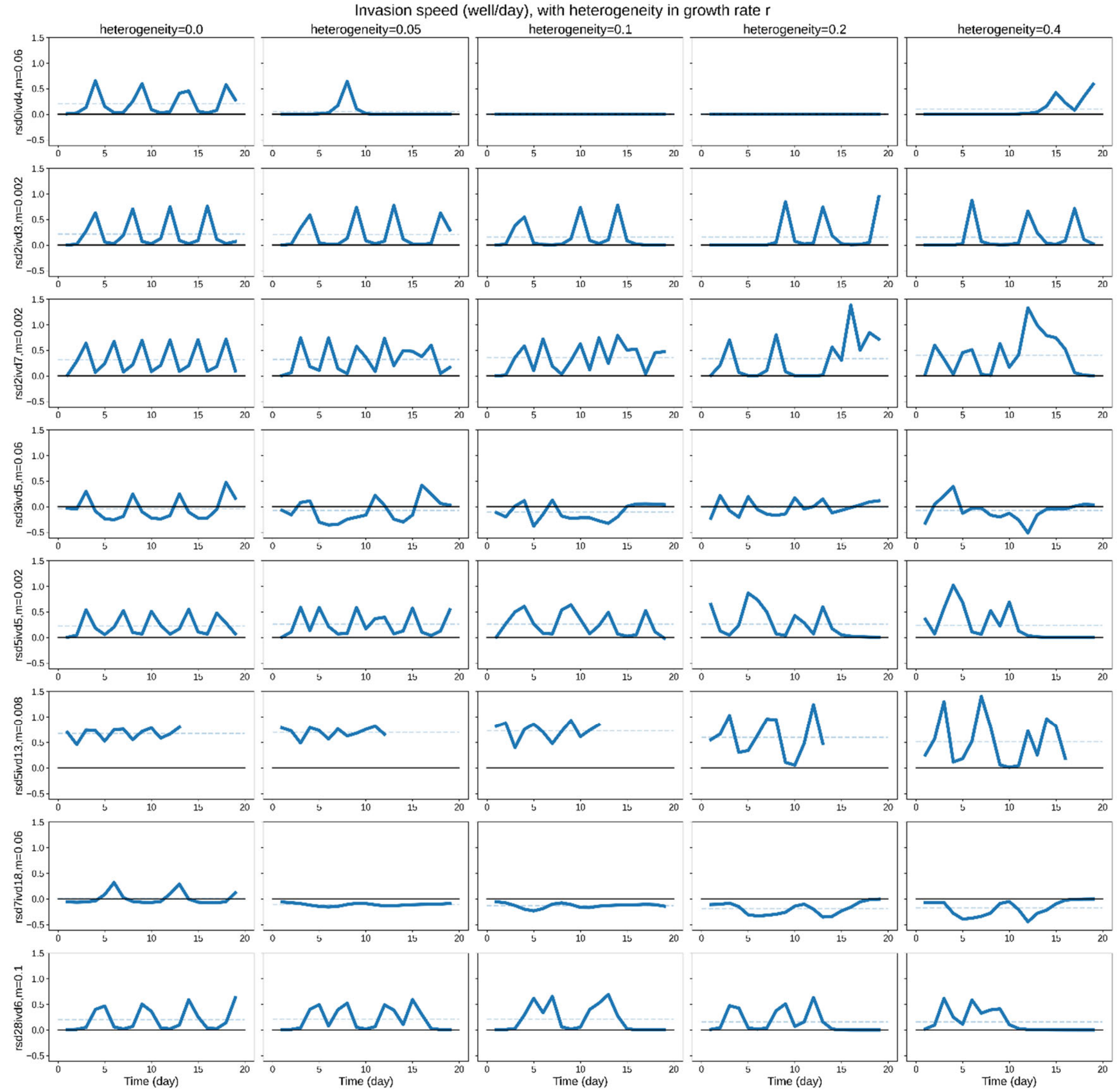

**Supplementary Figure 19. Daily invasion speed for invasion along patches with heterogeneous growth rate r.** Each row shows daily invasion speed (solid blue line) and mean invasion speed (horizontal blue dash) for an example with the given resident community (rsd), invader (ivd) and dispersal rate (m). The heterogeneity of growth rates r across patches increases from left to right. For given heterogeneity level H, the growth rate r for each species in each patch is multiplied by a random value drawn independently from the uniform distribution [1-H, 1+H], H=0 means no heterogeneity. To compare better with previous theoretical results, we defined the front position as the sum of invader fractions (divided by carrying capacity 1) when computing the invasion speed instead of the position above the 50% threshold; the conclusions stayed the same regardless.

### Supplementary Tables

#### Supplementary Table 1: Making Invasion Media (IM) from partial stocks

| **Partial stock components** | **$ddH_2O$** | | | | **pH** |
|---|---|---|---|---|---|
| **PS-C (carbon)** | 48.8mL | 0.5g soytone | 0.5g tryptone | 0.2g yeast extract | pH7 |
| **PS-A (acid)** | 47.4mL | Ammonia acetate (36.7mM 50mL* 77.0825g/mol * 10=1.414g) | Trisodium citrate·$2H_2O$ (8.2mM * 50mL * 294.10g/mol * 10=1.206g) | | pH6.88 |
| **10X PS-M (metals)** | 45mL | 4mL 1M $MgSO_4$ | 1mL 1M MnSO4 | | pH7 |
| **PS-P (phosphate)** | 49mL | $K_2HPO_4$ (0.5777M * 50mL * 174.18g/mol = 5.0312g) | $KH_2PO_4$ (0.4223M * 50mL * 136.09g/mol= 2.8735g) | | pH6.8 |
| **PS-G (20% w/w glucose)** | 40mL | 10g glucose | | | |
| **5X PS-S (salts)** | ~8.3mL (adjust to final volume of 10mL) | $(NH_4)Cl$ 0.8022g | NaCl 0.8760g | | add ~7µL 10N NaOH to make it pH7 |

PS-C must be freshly prepared and mixed with other components before each experiment. To prepare 50mL IM media: 5mL PS-C (1/10 complex carbon sources as in MRS), 500µL 10X PS-M (similar Mg and Mn as in MRS), 1mL PS-A (1/5 acid as in MRS), 1mL PS-G (1/5 glucose as in MRS), 400µL 5X PS-S (together with PS-A, make to similar ammonia and 1/2 salinity level as in MRS), 4mL variously diluted PS-P to reach different buffer concentrations (4mL undiluted PS-P makes 80mM final buffer concentration), and 38.1mL $ddH_20$.

**Supplementary Table 2: Liquid transfer for invasion experiment in two-strain system**

We use $x_n$ to denote well at position x, day n. Then $(x-1)_{n+1}$ is well at position x-1 (I.e. 1 well left to x), day n+1. The table below specifies liquid transferred into well $x_{n+1}$ from day n to day n+1. 1X and 1/100X refer to the undiluted and 1/100X diluted plate from the previous day.

| Dispersal rate m | Volume from direct dilution (nL) | Volume from dispersal of one side (nL) | Volume from $x_n$ (1X) (nL) | Volume from $x_n$ (1/100X) (nL) | Volume from $(x \pm 1)_n$ (1X) (nL) | Volume from $(x \pm 1)_n$ (1/100X) (nL) |
|---|---|---|---|---|---|---|
| 0.002 | 249.5 | 0.25 | 225 | 2450 | 0 | 25 |
| 0.008 | 248 | 1 | 225 | 2300 | 0 | 100 |
| 0.02 | 245 | 2.5 | 225 | 2000 | 0 | 250 |
| 0.038 | 240.5 | 4.75 | 225 | 1550 | 0 | 475 |
| 0.06 | 235 | 7.5 | 225 | 1000 | 0 | 750 |
| 0.1 | 225 | 12.5 | 225 | 0 | 0 | 1250 |
| 0.2 | 200 | 25 | 175 | 2500 | 25 | 0 |
| 0.4 | 150 | 50 | 125 | 2500 | 50 | 0 |

**Supplementary Table 3: Liquid transfer for invasion experiment in multi-strain system**

We use $x_n$ to denote well at position x, day n. Then $(x-1)_{n+1}$ is well at position x-1 (I.e. 1 well left to x), day n+1. The table below specifies liquid transferred into well $x_{n+1}$ from day n to day n+1. 1/10X refer to the 1/10X diluted plate from the previous day.

| **Dispersal rate m** | **Volume from direct dilution (nL)** | **Volume from dispersal of one side (nL)** | **Volume from x_n (1/10X) (nL)** | **Volume from x±1_n (1/10X) (nL)** |
|---|---|---|---|---|
| 0.02 | 245 | 2.5 | 2450 | 25 |
| 0.04 | 240 | 5 | 2400 | 50 |
| 0.06 | 235 | 7.5 | 2350 | 75 |
| 0.08 | 230 | 10 | 2300 | 100 |
| 0.1 | 225 | 12.5 | 2250 | 125 |
| 0.12 | 220 | 15 | 2200 | 150 |
| 0.22 | 195 | 27.5 | 1950 | 275 |
| 0.4 | 150 | 50 | 1500 | 500 |

**Supplementary Table 4: Liquid transfer for interaction experiment in two-strain system**

| | Lp% | Su% | Su_vol | Su_vol_ 1X | Su_vol_ 1/100X | Lp_vol | Lp_vol _1X | Lp_vol_ 1/100X |
|---|---|---|---|---|---|---|---|---|
| 1 | 100 | 0 | 0 | 0 | 0 | 250 | 225 | 2500 |
| 2 | 99.9 | 0.1 | 0.25 | 0 | 25 | 249.75 | 225 | 2475 |
| 3 | 99.6 | 0.4 | 1 | 0 | 100 | 249 | 225 | 2400 |
| 4 | 99 | 1 | 2.5 | 0 | 250 | 247.5 | 225 | 2250 |
| 5 | 98.1 | 1.9 | 4.75 | 0 | 475 | 245.25 | 225 | 2025 |
| 6 | 97 | 3 | 7.5 | 0 | 750 | 242.5 | 225 | 1750 |
| 7 | 95 | 5 | 12.5 | 0 | 1250 | 237.5 | 225 | 1250 |
| 8 | 93 | 7 | 17.5 | 0 | 1750 | 232.5 | 225 | 750 |
| 9 | 90 | 10 | 25 | 25 | 0 | 225 | 200 | 2500 |
| 10 | 87 | 13 | 32.5 | 25 | 750 | 217.5 | 200 | 1750 |
| 11 | 83 | 17 | 42.5 | 25 | 1750 | 207.5 | 200 | 750 |
| 12 | 80 | 20 | 50 | 50 | 0 | 200 | 175 | 2500 |
| 13 | 78 | 22 | 55 | 50 | 500 | 195 | 175 | 2000 |
| 14 | 75 | 25 | 62.5 | 50 | 1250 | 187.5 | 175 | 1250 |
| 15 | 72 | 28 | 70 | 50 | 2000 | 180 | 175 | 500 |
| 16 | 70 | 30 | 75 | 75 | 0 | 175 | 150 | 2500 |
| 17 | 68 | 32 | 80 | 75 | 500 | 170 | 150 | 2000 |
| 18 | 66 | 34 | 85 | 75 | 1000 | 165 | 150 | 1500 |
| 19 | 64 | 36 | 90 | 75 | 1500 | 160 | 150 | 1000 |
| 20 | 62 | 38 | 95 | 75 | 2000 | 155 | 150 | 500 |
| 21 | 60 | 40 | 100 | 100 | 0 | 150 | 125 | 2500 |
| 22 | 57 | 43 | 107.5 | 100 | 750 | 142.5 | 125 | 1750 |
| 23 | 53 | 47 | 117.5 | 100 | 1750 | 132.5 | 125 | 750 |
| 24 | 50 | 50 | 125 | 125 | 0 | 125 | 100 | 2500 |
| 25 | 45 | 55 | 137.5 | 125 | 1250 | 112.5 | 100 | 1250 |
| 26 | 40 | 60 | 150 | 125 | 2500 | 100 | 100 | 0 |
| 27 | 35 | 65 | 162.5 | 150 | 1250 | 87.5 | 75 | 1250 |
| 28 | 30 | 70 | 175 | 150 | 2500 | 75 | 75 | 0 |
| 29 | 25 | 75 | 187.5 | 175 | 1250 | 62.5 | 50 | 1250 |
| 30 | 20 | 80 | 200 | 175 | 2500 | 50 | 50 | 0 |
| 31 | 10 | 90 | 225 | 200 | 2500 | 25 | 25 | 0 |
| 32 | 0 | 100 | 250 | 225 | 2500 | 0 | 0 | 0 |

_vol: volume (μL) from precultures. _vol_1X, _vol_1/100X: volume (μL) transferred from 1X or 1/100X diluted plates, respectively.

**Supplementary Table 5: Liquid transfer for interaction experiment in multi-strain system**

The original experimental design consisted of 32 mixing fractions as in two-strain system, 25 of which used 1/10X diluted precultures as source plate as shown here. The other 7 mixing fractions (0.001 to 0.008 invader fraction by volume) used 1/100X diluted precultures as source plate, however, we encountered technical challenge as in highly diluted Pa cultures the bacteria gathered at surface, so these mixing fractions were discarded.

| | **CM%** | **Pa%** | **Pa_vol** | **Pa_vol _1/10x** | **CM_vol** | **CM_vol _1/10x** |
|---|---|---|---|---|---|---|
| 1 | 100 | 0 | 0 | 0 | 250 | 2500 |
| 2 | 99 | 1 | 2.5 | 25 | 247.5 | 2475 |
| 3 | 98 | 2 | 5 | 50 | 245 | 2450 |
| 4 | 97 | 3 | 7.5 | 75 | 242.5 | 2425 |
| 5 | 96 | 4 | 10 | 100 | 240 | 2400 |
| 6 | 95 | 5 | 12.5 | 125 | 237.5 | 2375 |
| 7 | 94 | 6 | 15 | 150 | 235 | 2350 |
| 8 | 93 | 7 | 17.5 | 175 | 232.5 | 2325 |
| 9 | 91 | 9 | 22.5 | 225 | 227.5 | 2275 |
| 10 | 89 | 11 | 27.5 | 275 | 222.5 | 2225 |
| 11 | 87 | 13 | 32.5 | 325 | 217.5 | 2175 |
| 12 | 85 | 15 | 37.5 | 375 | 212.5 | 2125 |
| 13 | 83 | 17 | 42.5 | 425 | 207.5 | 2075 |
| 14 | 80 | 20 | 50 | 500 | 200 | 2000 |
| 15 | 75 | 25 | 62.5 | 625 | 187.5 | 1875 |
| 16 | 70 | 30 | 75 | 750 | 175 | 1750 |
| 17 | 65 | 35 | 87.5 | 875 | 162.5 | 1625 |
| 18 | 60 | 40 | 100 | 1000 | 150 | 1500 |
| 19 | 55 | 45 | 112.5 | 1125 | 137.5 | 1375 |
| 20 | 50 | 50 | 125 | 1250 | 125 | 1250 |
| 21 | 40 | 60 | 150 | 1500 | 100 | 1000 |
| 22 | 30 | 70 | 175 | 1750 | 75 | 750 |
| 23 | 20 | 80 | 200 | 2000 | 50 | 500 |
| 24 | 10 | 90 | 225 | 2250 | 25 | 250 |
| 25 | 0 | 100 | 250 | 2500 | 0 | 0 |

CM: community. _vol: volume (μL) from precultures. _vol_1/10X: volume (μL) transferred from 1/10X diluted plate.

### Supplementary Texts

#### Supplementary Text 1: Parameter choice for mechanistic models

*Overview*

We used mechanistic models not to study the dynamics of the models themselves, but rather to assess the applicability and limitation of our parameter-free framework in diverse ecological scenarios. Therefore, we generally drew parameters from uniform distributions with rather wide ranges. We set one simulation time unit to resemble one hour in experiments, and chose the growth rate range to reflect the observed range in experiments. Specifically, we frequently observed doubling time between 1-4 hours in experiment, which gave rise to growth rates between 0.17 – 0.69; we adopted a similar range in mechanistic models. We set the range of other parameters accordingly based on growth rates, so that (1) their scales matched experimental intuition, (2) the majority of the community dynamics converged to a steady state as in experiment, (3) stable communities had high enough species diversity and thus complexity. We generally assigned invaders with higher growth rates and competition ability to increase the chance of a successful invasion, while making sure that such modifications didn't overrule other interactions. These considerations all together allowed for complex and diverse invasion scenarios, each driven by different species interactions and resulted in interaction curves of vastly different shapes.

*Generalized consumer-resource model*

The table below shows values for different parameters; brackets [] indicate the range for uniform distribution. For the list of all biochemicals, a fraction (fP) of biochemicals at the beginning of the biochemical list can be potentially consumed by species (i.e. resources), a fraction of resources (fC) at the end of the biochemical list can potentially inhibit species growth. A biochemical can be both a resource or a toxin, depending on the species. We allowed fP and fC to change in a wide range, which could potentially shift the type of interactions that drove invasion dynamics. On average, but not always, invaders as compared to residents have faster growth rates, can use more biochemicals, and are inhibited by fewer biochemicals. Besides, invaders can always use the 3 major biochemicals that are replenished when simulating daily dilutions.

| Parameter | Description | Values |
|---|---|---|
| nS | Number of species used to start a resident community (some of which are competitively excluded at equilibrium) | 50 |
| nR | Number of biochemicals | 30 |
| fP | Fraction of biochemicals that can be consumed by at least some species | [0.2, 0.8] for residents<br>[0.2, 1] for invaders |
| fC | Fraction of biochemicals that can inhibit the growth of at least some species | [0.2, 0.8] for residents<br>[0, 0.8] for invaders |

| | | |
|---|---|---|
| $r_i$ | Innate growth rate of species i | [0.1, 0.7] for residents<br>[0.6, 0.8] for invaders |
| $p_{ij}$ | Consuming preference of species i on biochemical j | For the 3 biochemicals that serve as main resource:<br>$\begin{cases} [0,1], P = 0.2 \\ 0, P = 0.8 \end{cases}$<br>for invader 0.1 is added to the drawn value.<br>For all other biochemicals:<br>$\begin{cases} [0,1], P = 0.5 \\ 0, P = 0.5 \end{cases}$ |
| $b_i^{max}$ | Maximum secretion rate of species i | [0.2, 1] |
| $b_{ij}$ | Relative rate of species i secreting biochemical j | $\begin{cases} [0,1], P = 1/3 \\ 0, P = 2/3 \end{cases}$<br>After random drawn from the above distribution, normalize for each species i so that $\sum_j b_{ij} = b_i^{max}$ |
| $c_{ij}$ | Inhibition coefficient of species i by biochemical j | $\begin{cases} [0,20], P = 1/3 \\ 0, P = 2/3 \end{cases}$ |

*Generalized Lotka-Volterra model*

The table below shows values for different parameters; brackets [] indicate the range for uniform distribution. The interaction coefficient matrix consists of weak, strong, and no interactions. The ratio of these three types was adjusted to allow strong interactions while still maintaining diverse and stable resident communities. On average, invaders grow faster than residents and inhibit residents more strongly. We first generated a species pool of 1000 resident species and 100 invaders, including growth rates and pairwise interaction coefficients. From this pool, we randomly sampled nS=50 residents and 1 invader for each invader-resident pair, together with the previously generated parameters.

| **Parameter** | **Description** | **Values** |
|---|---|---|
| nS | Number of species used to start a resident community (some of which are competitively excluded at equilibrium) | 50 |
| $r_i$ | Innate growth rate of species i | [0.1, 0.5] for residents<br>[0.45, 0.6] for invader |
| $A_{ij}$ | Interaction coefficient of species j on species i. Positive values indicate facilitation. | If both species are residents:<br>$\begin{cases} [-0.5, 0.1], P = 0.2 \\ [-2, 0], P = 0.4 \\ 0, P = 0.4 \end{cases}$<br>If species i is resident and species j is invader: [-3, 0]<br>The generated interaction matrix is further modified by setting diagonal terms $A_{ii}$ to -1 for self-inhibition. |
